# Accurate prediction of CRISPR editing outcomes in somatic cell lines and zygote with few-shot learning

**DOI:** 10.1101/2024.11.08.622621

**Authors:** Weizhong Zheng, Lu Yu, Guoliang Wang, Shaoxian Cao, Kenso Ho, Jun Song, Chen Cheng, Joshua W.K. Ho, Xueqing Liu, Meng Wu, Zhonghua Liu, Huili Wang, Pentao Liu, Guocheng Lan, Yuanhua Huang

**Author notes:** Correspondence, {, }. co-first authors (Weizhong Zheng, Lu Yu).

## Abstract

The CRISPR-Cas system has revolutionized gene editing, while its outcome prediction remains unsatisfactory, especially in new cell states due to their distinct DNA repair preferences. In this study, we introduce inDecay, a flexible system for predicting CRISPR editing outcomes from target sequence, returning probabilities of nearly the full spectrum of indel events. Uniquely, inDecay utilizes informative and parameter-efficient features for each indel event and incorporates cell-type-specific repair preferences through a multi-stage design. While both inDecay and existing methods achieve accurate results for prediction within cell lines, only inDecay with transfer learning can retain the high performance for cross-cell line prediction. We then applied inDecay to mouse embryo editing using our newly generated data and observed remarkable accuracy by including as few as 30 fine-tuning embryonic samples. Notably, inDecay is the first software to predict embryonic editing. Therefore, our few-shot learning-supported system may accelerate guide RNA prioritization in mouse model generation, mammal embryonic gene editing, and cellular therapeutics.

## Introduction

The CRISPR-Cas system enables precise and efficient genome editing (Jinek et al., 2012; Cong et al., 2013; Ran et al., 2013). A wide range of its derived applications, including generating model animals and treating genetic diseases by correcting pathogenic genetic variations (Polak et al., 2015; Blanpain et al., 2011; Bowling et al., 2020), require precise control of potential editing outcomes. To this end, it is helpful to have an unbiased and accurate prediction of the repair outcomes during guide RNA (gRNA) design to avoid undesired editing outcomes.

CRISPR-induced genome editing primarily occurs through the repair of a double-stranded break (DSB) at the site targeted by the guide RNA (Eker et al., 2009; Ceccaldi et al., 2016), producing a diverse array of repair products. For template-free CRISPR editing, the main pathways involved are classical Non-Homologous End Joining (cNHEJ), which produces small insertions and deletions, and Microhomology-Mediated End Joining (MMEJ), which generally produces larger deletions with the existence of short homologous sequences (Liu et al., 2021; Xue and Greene, 2021). The choice of the repair pathway and the randomness within each repair process make up a complicated repair profile. The repair profile is regulated at two levels. Globally, the expression of DNA repair genes shapes the overall preference over different repair pathways (Bothmer et al., 2017; van Overbeek et al., 2016). Locally, the sequence context flanking the cleavage site determines the probability of each potential insertion or deletion (indel) based on the preference of indels by different repair pathways.

Recently, highly scalable self-targeting editing techniques have emerged, making it possible to comprehensively profile CRISPR-induced DNA repair events for a target sequence (Shen et al., 2018; Allen et al., 2019; Chen et al., 2019). The generated data offer unique opportunities to decipher the impact of local sequence context on repair patterns and were used to develop computational methods for predicting CRISPR editing outcomes, with prominent examples including inDelphi (Shen et al., 2018), ForeCasT (Allen et al., 2019) and Lindel (Chen et al., 2019). However, existing methods have primarily been trained on datasets measured in a certain cell line (or pooling multiple lines) and learned the mapping from sequence to repair profile, ignoring the variability of the global environment between cell types or conditions (Fuchs and Blau, 2020; da Silva et al., 2021). Consequently, their predictions may be inaccurate for cell types that exhibit distinct preferences for repair pathways (Shen et al., 2018; Allen et al., 2019).

Embryonic cells exhibit distinct DNA repair activity compared to somatic cells, particularly during early developmental stages (Khokhlova et al., 2020). Studies suggested that embryonic cells may favor the NHEJ repair pathway at certain stages, possibly due to the absence of specific proteins and shifts in cell cycle phase distribution (Tichy and Stambrook, 2008; Khokhlova et al., 2020; Morozumi et al., 2024). As a result, predicting CRISPR editing outcomes in embryonic cells poses greater challenges for current methods. For example, although inDelphi has developed variants tailored to several cell types (Shen et al., 2018), its predictions tend to over-represent microhomology-mediated end joining (MMEJ) repair outcomes. Furthermore, these methods have shown poor performance when applied to germline editing (Naert et al., 2021, 2020).

In this study, we present *inDecay*, a multi-stage deep learning method for predicting the CRISPR-induced editing outcome given the target sequence. Our method particularly considers the variability of the DNA repair system across different cell types or conditions by explicitly modeling the ratios of different repair pathways (insert, MH deletion, and NH deletion). Moreover, we designed an informative encoding scheme for each deletion event based on its deletion length, MH track, and deletion start site. The encoded deletion features enable efficient estimation of the probabilities of deletion events and allow for post hoc customization of the model using user-adjustable hyperparameters. Our results demonstrate the superiority of *inDecay* over existing methods on multiple cell lines. We particularly emphasize the use of few-shot learning, which effectively facilitates the generalization of our model to new cell lines. Moreover, we demonstrated that this capability of transfer learning plays a critical role in accurate prediction in mouse embryos in our newly generated *in vivo* datasets. Therefore, with its flexibility and accuracy, *inDecay* holds great potential for broader application in CRISPR-based genome editing studies.

## Results

### Highly informative event-based features enable a flexible and lightweight model for repair outcome prediction

The DNA repair outcomes of the CRISPR-induced double-strand break depend on both the sequence context of the target site and the cellular environment as defined by the relative activities of different repair pathways (Bothmer et al., 2017; van Overbeek et al., 2016). We designed a novel repair outcome prediction method inDecay that can consider the impact of both layers. Our method inDecay first captured the sequence context by detecting all possible microhomologous (MH) sequences (referred to as MH-track later) within the target sequences. This allowed us to quickly identify the MH deletion events and craft features for each event. Next, the preference for different repair outcomes was captured by a multi-stage learning system in a cell type-specific manner, which used manually crafted features to predict the probability of each event (indel).

The strength of MMEJ events is largely decided by the length and the starting position of the MH-tracks. A long MH-track that is close to the cut site can sometimes result in a single MH deletion event dominating the entire repair profile (Brinkman et al., 2018). Characterizing MH tracks and annotating the MH deletion from hundreds of possible deletion events is thus important. To quickly identify all the MH-tracks in the target sequence, we converted the problem of searching all possible homologous sub-sequences into a diagonal-line detection problem. This was achieved by matching the left and right cleaved fragments into a two-dimensional matrix similar to the sequence substitution matrix (Figure 1A), which was traditionally used to score aligning nucleotides. Then, MH-tracks can be efficiently identified by applying convolution filters onto the matrix (see Method), avoiding iterative string matching by previous methods (Allen et al., 2019; Chen et al., 2019). The identified MH-tracks were then used to annotate the corresponding MH-deletions in terms of their deletion start site, maximum MH length, and deletion length.

**Figure 1.**
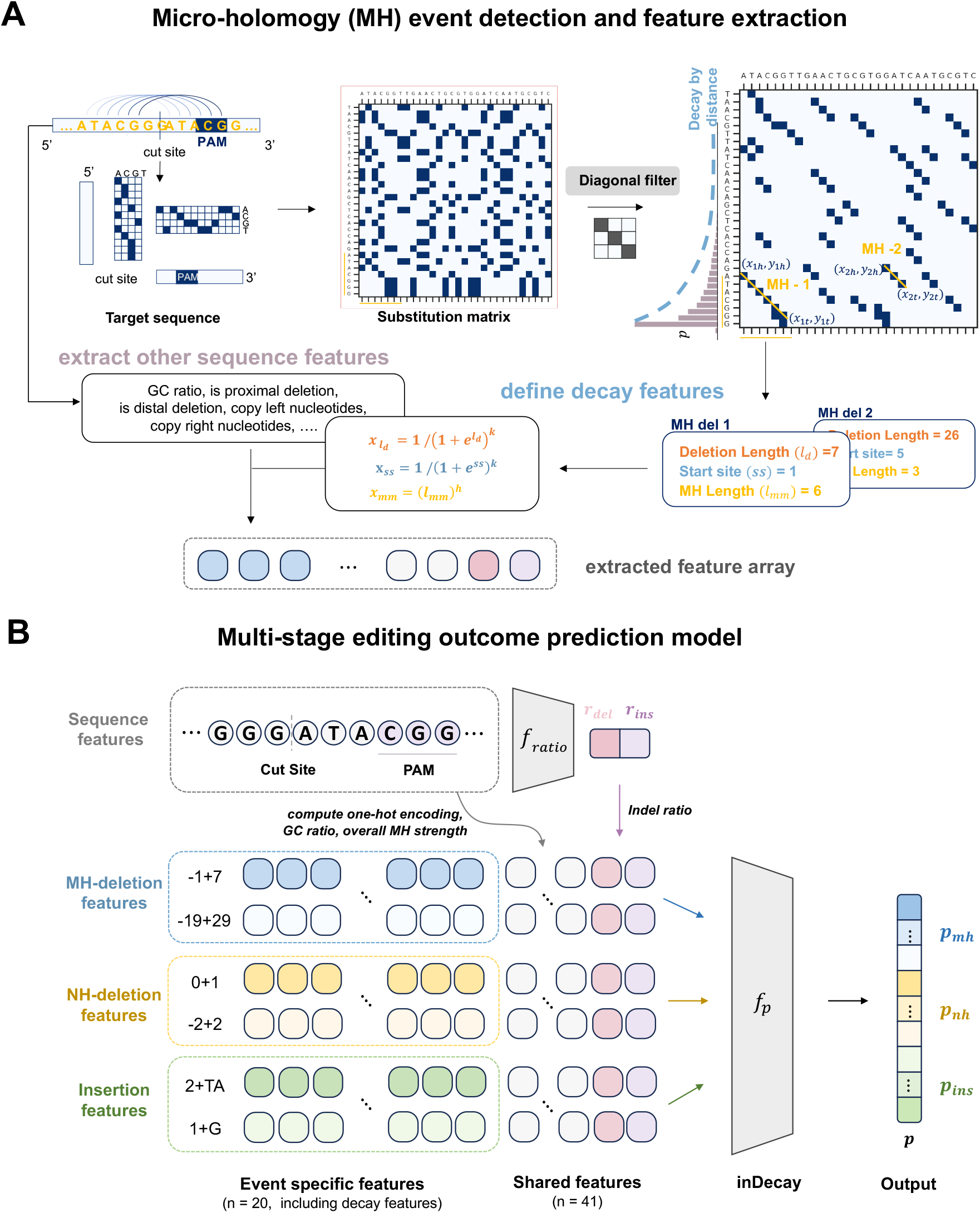
Schema of the inDecay model. **A** The workflow of MH event detection. A target sequence was first cut into two fragments at the cut-site and then computed the pair-wise nucleotide substitution matrix. The y-axis (first dimension) of the matrix represents the left fragment, and the x-axis represents the right fragment. The filtered matrix has only diagonal lines left. The y-coordinate of the tail end indicates the start site of the lines. The deletion length was the addition of the y-coordinate with the x-coordinate of the tail-end. Features were extracted based on the detected deletion events and acted as the input for the multi-stage model. **B** The multi-stage prediction models. After identifying the MH deletion events for a given target sequence, the full event frequency was predicted via the *f*_*p*_ module, which transforms the features of each event into frequency. The whole feature set comprises 20 features tailored to each indel event based on their categories, along with 36 one-hot encoded sequence features shared across all events and the GC ratio, the overall MH-strength, and the ratios of insertion and deletion predicted by the *f*_*ratio*_ module.

We observed a universal frequency decay of deletion events with respect to the deletion length as well as the start site (proximity of starting position to the cut site) across different cell types (see Figure 1A). However, depending on the sequence context, this pattern can be violated due to larger deletions that were induced by long MH-tracks. We therefore designed features in a first-principle manner to represent this statistical frequency decay using the deletion start site, maximum MH length, and deletion length extracted by the method mentioned above. The deletion length and start site were transformed by a decay function and the MH length was regulated by a power function (Figure 1A and Methods), with each term having its own hyper-parameter.

In the next stage, we utilized the annotated events and first-principle-derived features to predict the full spectrum of repair outcomes along with additional sequence features. The prediction was done via two steps. The first step involved the ratio module of inDecay to predict the overall proportion of deletion and insertion from the 9-bp nucleotides before the PAM site. This proportion was highly predictable using a linear model. In fact, different studies have reported a consistent nucleotide usage when it differentiates deletion-prompt and insertion-prompt sequences (Shen et al., 2018; Allen et al., 2019; Chen et al., 2019).

In the second step, more features were generated based on the category, as well as the deleted or inserted nucleotides of each indel. Following that, the inDecay frequency module would transform the finalized feature array of an event into its probability using a neural network. These event-specific features were constructed differently for the three classes, including microhomology (MH) deletions, non-MH deletions (NH), and insertion. NH deletion features were similar to MH deletion features with several features setting to be zero. Insertion features mainly captured the indel size and the complement of inserted nucleotides to the flanking sequences. Additionally, we added three types of features shared among all events to inform the overall property of the target sequence, including the insert-del proportion from the first stage, the overall MH-strength derived from the filtered substitution matrix, and the one-hot encoded 9-bp sequences close to the PAM site (36 features). The shared features reduced the inductive bias of the neural network by learning hidden variables that can complement the hand-crafted features. Taken together, we curated 61 information-dense features as the input and we were therefore able to use a light-weight neural network with only two hidden layers to accurately predict the full probability.

In summary, we proposed an event-based multi-stage learning method inDecay to predict the CRISPR editing outcome. A new MH-track search algorithm was integrated into inDecay to quickly emphasize the important MH-deletion features. inDecay combined feature engineering that was obtained from first-principle knowledge and automatic feature learning by neural networks, making it a light-weight yet powerul model.

### inDecay accurately predicts the repair profiles for cell lines

To evaluate the performance of inDecay for different cellular environments, we applied it to a multi-cell line dataset generated by Allen & Crepaldi et al (Allen et al., 2019), which profiled the repair outcomes of over 35,000 guide RNAs for all 5 cell lines (Figure S1). Among them, we selected 1133 highly covered guide RNAs containing over 1,000 total counts across cell lines as the test set and kept the remaining samples that had over 500 counts to train our inDecay model. The repair profiles of the targeted sequences have manifested a strong variability among cell types in several aspects (see Figure S2). Baseline models Lindel and FORECasT provided only a single model without cell type specification. To ensure a fair comparison, we retrained Lindel and FORECasT for each cell line, using the same training and test split. Another baseline method inDelphi has trained different versions on an independent dataset for 4 different cell lines. Among them, mESC and K562 overlapped with our training data set. Due to the difficulties of reproducing inDelphi’s pipeline, we only compared inDelphi on mESC and K562 data. We then benchmarked inDecay with the baselines on the held-out test set. To thoroughly assess the predicted outcome distribution of each method, a variety of evaluation metrics were applied, including both those previously utilized and newly developed ones (see Methods). FORECasT’s datasets involved two experimental replicates for each cell line, repair profiles from two replications were used to determine the upper bound of each evaluation metric.

We began by evaluating the yield of the most frequent events through the recall of top-5 predicted indel size and events. CRISPR editing typically produces a long-tail distribution of indels. Although each target sequence can generate over 300 unique indels, the top 5 accounted for 65.1% of the total (averaged over 59,337 samples from 5 cell lines, Figure S3). Adding five more indels increased this cumulative frequency to only 77.2%, indicating that the first five indels were the dominant events. Here, inDecay showed the highest top-5 recall in terms of indel sizes and detailed events (3.6 for inDecay in Figure 2A, 3.3 in 2B). inDecay can largely outperform Lindel and was better than FORECasT and inDelphi. Consistently, inDecay also achieved the highest top-10 recall (Figure S4A). These together suggested that inDecay can robustly reveal the most dominant and sub-major indels despite using much fewer features than previous methods.

**Figure 2.**
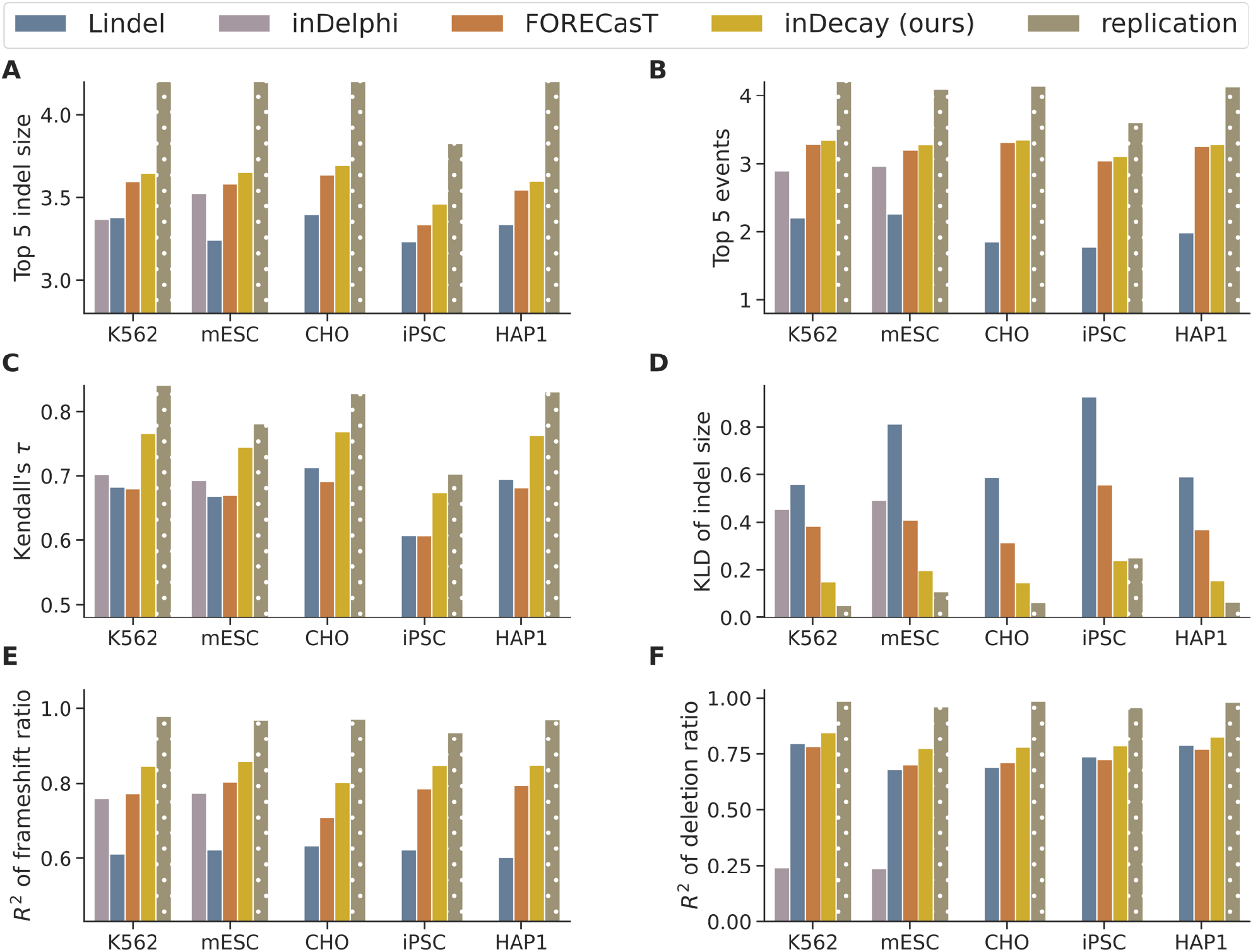
Performance of inDecay on FORECasT’s multi-cell type datasets. Evaluation of model performance across different cell lines using various metrics. Models, including Lindel, inDelphi, FORECasT, and inDecay (ours), were trained on the same dataset and tested on 1,133 guides. inDelphi was evaluated only for mESC and K562. The X-axis showed the cell type and the y-axis showed the evaluation metrics for all the subplots. The color of the bar denotes different methods: replication (dots) was calculated from the repair profile of the two experimental repeats. **(A-B)** Event level top 5 indel length and top 5 events recall calculated between predicted and observed event probability. The value of top-k recall directly indicated the number of overlapped events with the true top-k. i.e., *top*5 = 3 means 3 out of 5 most highly predicted indels are truly the top-5 indels. **(C-D)** Kendall’s rank correlation coefficients and KL divergence for measuring the overall similarity between the predicted and observed distribution. **(E-F)** The accuracy of the model recapitulates the proportion of different types of indels, the proportion of Frameshift ratio, and the proportion of deletion events in the distribution.

Next, we looked into the prediction over indel length by grouping events according to the editing length. In CRISPR/Cas gene editing experiments, the length of the editing outcome is essential for evaluating editing efficiency, specificity, and functional phenotype. By collapsing repair events of the same length, we transformed highly sparse event probability into a dense indel-length distribution consisting of 41 elements. The transformation allows for accurate measurement of the reconstructed distribution as it avoids the bias caused by the null values of the rare indels (as opposed to the Mean Squared Error used in (Chen et al., 2019)). When evaluating the overall similarity of indel length, inDecay could present a KL divergence around 10% and 32% lower than inDelphi and the FORECasT (Figure 2D). We also included Kendall’s *τ* coefficient, which considered the ordering within 41 indel lengths. inDecay could predict the most concordant ranking of indel length, with a mean Kendall’s *τ* of 0.7 and 0.73, respectively (Figure 2C). Interestingly, while we found FORECasT achieved comparably well top-5 recall, its Kendall’s *τ* (among 41 indel lengths) was substantially lower than that of inDecay, indicating its suboptimal in predicting the ranking of top events (similarly in the KL divergence metric). These results together demonstrated inDecay’s ability to accurately recapitulate the indel length distribution at different scopes.

We subsequently inspected the model predictions concerning the proportion of different categories of repair events. The frameshift ratio *R*^2^ (FSR2) indicates the probability of silencing a gene with one edit. Across all cell lines, inDecay demonstrated the highest frameshift ratio *R*^2^. Specifically for mESC cell, inDecay reached an FSR2 of 0.859 with a large improvement over existing baselines (0.622 for Lindel, 0.77 for inDelphi, and 0.804 for FORECasT), thus improving our ability to estimate the efficiency of introducing loss-of-function indels (Figure 2E & Figure S4). Additionally, we investigated the predicted ratio of deletion events to insertion events (Figure 2F). Methods adopting a multi-stage design showed an advantage over the end-to-end baseline FORECasT (Figure 2F). Lindel and inDecay were among the top performers. Unexpectedly, inDelphi displayed disproportionally low *R*^2^, attributed to a general overestimation of the deletion events (Figure S4B).

In summary, our method inDecay could accurately predict the repair profiles for different cell lines. Combining the advantage of an event-based model and multi-stage design, inDecay could make predictions resembling the actual event probability regarding the most frequent events, the distribution of indel length, and the proportion of different indel types.

### Transfer learning enables effective cross-cell line prediction for inDecay

In practice, CRISPR-Cas genome editing could be carried out by researchers across a diverse array of cell types and organisms, many of which were not encompassed within the dataset employed by our model. Transfer learning represents a way to effectively overcome the cell type variability for models pre-trained on another cell line (3A). To demonstrate this, we selected two cell pairs, CHO-mESC and CHO-iPSC, based on their KL divergence in indel length distribution (see Figure S2). To assess the minimum number of samples required to overcome cell line variability, we performed multiple transfer learning experiments where the source model, pre-trained on CHO cells, was fine-tuned with increasing sample sizes of the target data set. Specifically, we used 10, 30, 50, 100, 300, 500, and 1000 guides for inDecay pretrained on CHO cells to finetune the models for mECS and iPSC cells. The finetuning data were randomly sampled from training-set guides with over 500 read counts, and the transferred models were evaluated with 1133 test-set guides from the target cell line.

When directly applied to a different cell line without fine-tuning, inDecay showed a significant decrease in multiple metrics compared to models that were trained from scratch on the corresponding cell line. For example, the frameshift ratio R2 of inDecay has both dropped by around 0.3 in mESC and iPSC. However, the gap between CHO and mESC can quickly covered with merely 30 samples (mean FSR2 =0.74 and 0.69, Figure 3B, C). Other metrics also experienced the greatest marginal improvement when finetuning with 30 samples, including Top 5 indel length, Top 10 indel length, and KL divergence. The performance gain conveyed by finetuning became almost saturated for inDecay with more than 50 samples (Figure 3C). Although the metrics for Top 5 event recall and Top 10 event recall decreased after fine-tuning with 30 samples, we analyzed the ranking order of the Top 5 and Top 10 events using the Kendall-tau coefficient (Top 5 events tau and Top 10 events tau). We found that after fine-tuning with 30 samples, the order of the predicted top events showed significant improvements compared to the baseline. For more distal cell line iPSC, the improvement above 30-50 samples also became marginal. It took around 300 samples for a stationary performance, probably due to the larger gap in repairing preference. While finetuned inDecay also benefited from seeing more samples, the best performance was generally not on par with inDecay.

**Figure 3.**
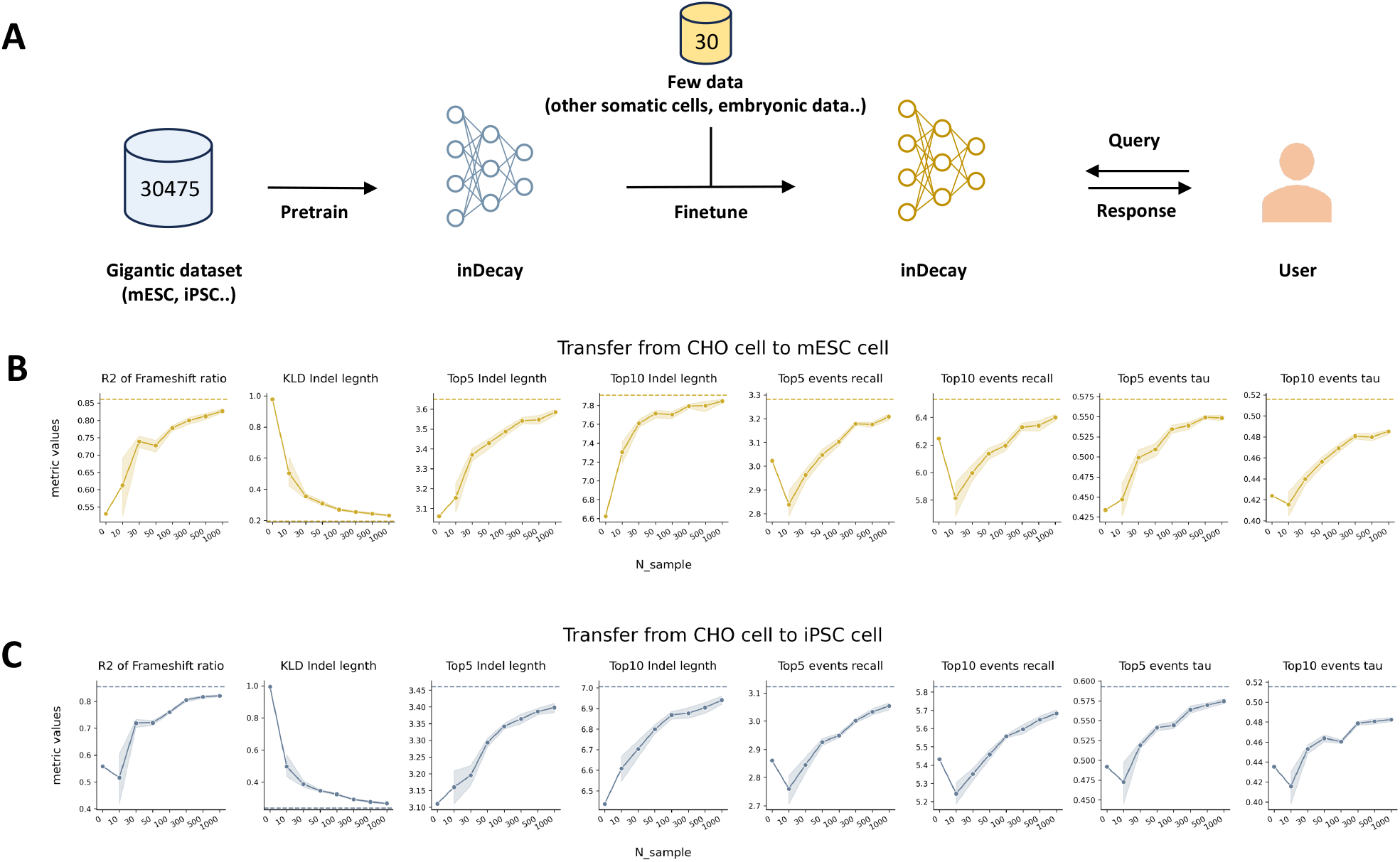
Transfer inDecay to another cell type with different sample sizes. **(A)** the workflow of inDecay. **(B-C)** Evaluating the transferred models on test set (n=1133) of mESC **B** and iPSC **(C)**. The gray band of the line plot shows the value range of 10 training repeats, the dots were the average performance over the repeats, and the dashed lines were the benchmark Performance of inDecay on FORECasT’s mESC (yellow) and iPSC (blue) datasets.

In summary, inDecay could gradually catch up with the repairing preference of another cell by fine-tuning with more data. Especially, inDecay relied on merely 30 to 50 samples to reach a satisfying similarity to the target cell line. And taking advantage of inDecay, users can quickly customize the model for unseen somatic cells or even embryos with very few fine-tuning samples (3A).

### inDecay can quickly adapt to embryo editing patterns using few-shot learning

An important application of CRISPR-Cas template-free editing in biology is to produce gene-knockout animal models. Given the limitations in the availability of mouse embryos, it is crucial to ensure effective editing outcomes to maximize the yield of knockout embryos. However, embryo editing normally involves microinjection or electroporation to deliver CRISPR-gRNA complex, and embryonic cells hold unique repair preferences, making the editing profile of embryos distinct from other cell lines. Previously established prediction systems, tailored for self-targeting data, have been found inaccurate for embryo editing (Naert et al., 2021, 2020).

To address the challenge of predicting embryonic editing outcomes, we designed 30 gRNAs targeting 22 genes and repeated the CRISPR-editing in a large number of mouse embryos for each gRNA (Figure 4A). The targeted regions were PCR amplified and sent for Sanger sequencing. The sequencing results underwent preprocessing and filtering to obtain the editing profile of the gRNA for the gene (see Methods). Consequently, we compiled a dataset of embryonic editing outcomes targeting 30 different genomic loci, collected from a total of 981 mouse embryos. As our methods have demonstrated a sample efficient transferring ability, we employed few-shot learning on inDecay to capture the patterns of embryonic editing. To facilitate few-shot learning and ensure fair evaluation with limited data, we applied leave-one-out cross-validation to split the fine-tuning and testing samples. In each fold, we evaluated the prediction results of one guide while using the remaining 29 guides for training and validation. By aggregating the results from all 30 folds, we obtained the model performance for all 30 guides.

**Figure 4.**
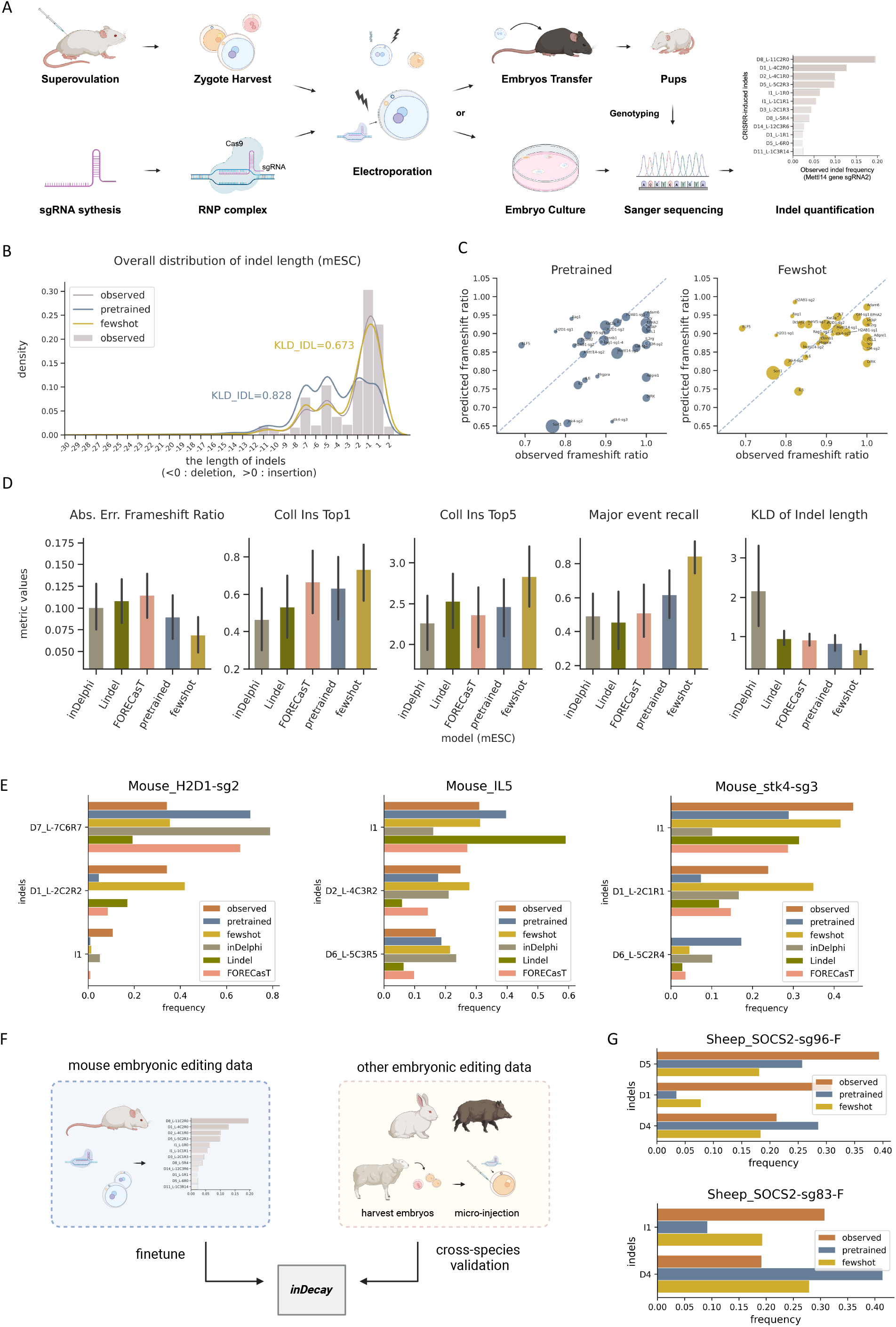
Transferring and evaluating inDecay in newly generated embryonic editing dataset in mouse and mammals. (Caption next page.) Figure 4. (Last page.) **(A)** The illustration of the workflow of collecting mouse embryonic editing data. **(B)** Overall indel length distribution of the 30 guide RNAs on mouse embryos (purple bar) and their predicted distribution by inDecay only pre-trained in somatic cell lines (pre-trained, blue curve) and fine-tuned on our zygote data (fewshot, yellow curve). Kullback–Leibler divergence value (KLD_IDL) was computed between the observed and each of the predicted indel length distributions (lower KLD_IDL value is better). **(C)** Scatter plot compared the observed frameshift ratio (x-axis) with the predicted frameshift ratio per guide on the pre-trained model (left) and finetuned model (right), where the averaged error was shown in panel (D). **(D)** Performance of pre-trained and few-shot on different metrics. From left to right shows, the absolute error of frameshift ratio (|predicted ratio – observed ratio|), top1 & top5 event recall collapsing insertion events of the same length, major events (*β* > 0.20) recall ratio, KL divergence of indel length distribution. **(E)** Mouse sample examples showing the most frequent events of observation (observed, orange) and their predicted frequency by pre-trained model (pre-trained, blue), finetuned model (few-shot, yellow), inDelphi of version mESC (inDelphi, grey), Lindel (Lindel, olive), and FORECasT (FORECasT, salmon). Indels were labeled by identifiers, a CIGAR-like string. The former part suggests whether an event is Insertion or Deletion and the length of the indel. The latter part indicates the editing start site L and end site R relative to the cut site as well as the number of nucleotides matching the inserted / deleted sequence suggested by the number following C. **(F)** The illustration of the workflow of collecting other species’ zygote editing data for cross-species validation. **(G)** Sheep sample examples showing the most frequent indel length events of observation (observed, orange) and their predicted frequency by pretrained (pretrained, blue) and finetuned model (fewshot, yellow).

**Table 1.**
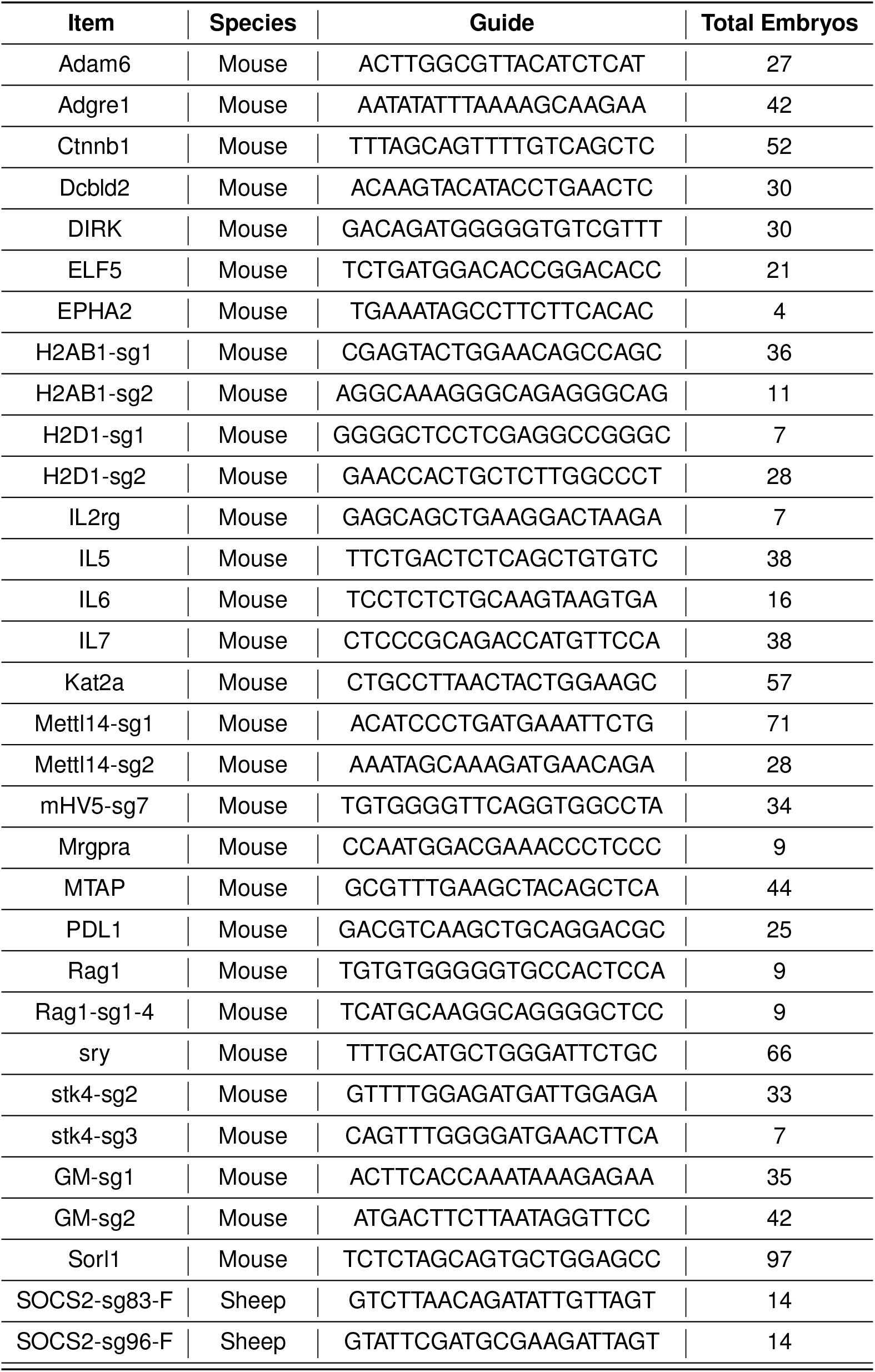
The guide sequence of designed gRNA and total embryo samples used.

Overall, our pre-trained inDecay model outperforms current mainstream tools in embryonic editing prediction, with our few-shot learning model demonstrating even superior performance (Figure 4B, C, D). The improvement gains from few-shot fine-tuning was consistent across different source cell lines (Figure S6). In mouse zygote editing, peak activities were observed for small indels (1-bp deletion, 1-bp insertion), as illustrated in Figure 4B, where the yellow curve of our finetuned inDecay model accurately captured this trend, with detailed KL divergence of indel length shown in the last panel in Figure 4D. However, the distribution captured on mESC cells showed additional peaks at 5 and 7 deletions, suggesting that embryo editing data indeed possesses unique repair preferences that cannot be revealed by self-targeting data. The scatter plot in Figure 4C showed the comparison between observed and predicted frameshift ratios. The pre-trained model tended to underestimate the frameshift ratio, while after fine-tuning, the frameshift predictions aligned more closely with the observed data to guide the gene knockout rate (around the diagonal line), except for some gRNAs with smaller embryo sample sizes (smaller point size) which may not capture the real editing patterns enough.

In Figure 4D, we compared our pre-trained model and few-shot model with other mainstream tools, demonstrating that our few-shot models significantly outperformed the pre-trained models and other tools across all metrics. Our few-shot model not only reduced the absolute error between predicted and observed frameshift ratios but also exhibited significant improvement in aggregated metric Coll Ins Top1 and Coll Ins Top5, respectively denoting the Top1 and Top5 events recalls merging insertion events. These metrics were introduced was because the pipeline used for parsing the editing products from Sanger trace data cannot recover the precise inserted nucleotides, which was different from the data we previously resolved from the next-generation sequencing. Notably, our few-shot learning model recalled an average of 84.5% major events (events with a ratio no less than 20% for each gRNA in the observed data can also be predicted as no less than 20% events) among the gRNAs, while inDelphi, Lindel, and FORECasT with only around half of its performance. These above demonstrated few-shot fine-tuned model performed the best not only in capturing the dominated event but also recalling the nearly full spectrum of indel events with correct order. Diving into the events prediction for each gRNA prediction, the precision and accuracy of our few-shot model were further highlighted in the predictions per gRNA in Figure 4E where the listed events were the union of Top2 events predictions among all tools.

Furthermore, the fine-tuned few-shot model was validated in zygotes of other species, as shown in Figure 4F. Notably, even without fine-tuning on the targeted species’ embryonic data, our few-shot learning model trained on mouse embryonic dataset led to better indel event predictions than the naive model pre-trained on somatic cell lines, showcasing it indeed captured more embryonic editing patterns (Figure 4G).

## Discussion

The CRISPR-induced DSB repair procedure displays substantial cell type variability. In this study, our re-analysis of FORECasT’s datasets showed that the difference in MH-strength and deletion-insertion ratio can partially explain such variability. To construct a versatile prediction system that can capture the intricate variations across different cell lines, we introduced inDecay. inDecay involves a multi-task ratio predictor to capture the cell line-specific ratio preference. Coupled with our unique MH-event detection system, inDecay featureed each MMEJ and c-NHEJ deletion event with several highly informative decay terms, which allowed it to predict the event probability in a highly parameter-efficient manner. Through extensive benchmarking with inDelphi, Lindel, and FORECasT, we showed that inDecay can outperform other state-of-the-art algorithms and can predict repair outcomes with multiple characteristics aligned with the true observation (see Figure 2). The efficient design of inDecay allowed it to be adopted to another cell line with as few as 30 to 50 samples (see Figure 3).

Embryo editing shows a highly distinct repair preference compared to somatic cells, yet it represents an important application of CRISPR editing. Current repair outcome prediction methods have not been thoroughly evaluated in the embryo system. In this study, we collected DNA repair profiles from more than thirty genomic sites across thousands of mouse zygotes. We found that even methods trained on mESC cell lines could not accurately recover the observed editing outcomes in mouse embryos (see Figure 3D). Only after fine-tuning the model on our mouse dataset could inDecay perform well on unseen samples and generalize to predict the editing in sheep embryos.

One hurdle in developing a generalizable editing outcome prediction system lies in the under-characterization of cell types and species-level variability of repair outcome preferences. As early efforts, both FORECasT and inDelphi have employed the self-targeting system in the context of several different cell lines. However, it remains unclear how representative these cell lines are for their species and their counterparts in the living organism. While it seems challenging to conduct large-scale self-targeting assays in diverse cell types, the gene expression profiles of different cells have mainly been sequenced and become easily accessible. Linking the gene expression to inform the repair preference of uncharacterized cells might stand as a promising path. Recent technical advances by repair-seq (Hussmann et al., 2021) have demonstrated how the repair outcomes vary along different perturbed expression contexts, revealing the genetic dependence and refined heterogeneity in DSB repair pathways. We can leverage inDecay’s predicted event distribution as a prior. Given the activities of the core DSB repair pathways of a cell, we can reweight this prior according to the pathway to obtain the final repair outcome for an unseen cell type.

Large DNA deletion is a type of unintended repair outcome not typically presented by self-targeting systems. In our mouse embryo data, we observed large deletions ranging from hundreds to thousands of base pairs, extending far beyond the target site. Contradictory arguments have been reported regarding its relationship with the MMEJ pathway (Owens et al., 2019; Kosicki et al., 2022). A very recent study suggested that additional repair mechanisms were involved in producing large deletions, rather than solely relying on the MMEJ and c-NHEJ pathways (Hwang et al., 2024). Predicting the emergence of large deletions from the sequence alone remains challenging and requires further investigation.

As CRISPR-based genetic therapy is getting ready for clinical use, it would be necessary to have an estimation of the precise editing efficiency ahead of the therapy. Considering the security of the individual’s genetic data and other ethical issues, the repair outcome data on the actual human embryos or stem cells is foreseeable to be very limited. A flexible and parameter-efficient prediction system like inDecay will largely benefit the few-shot transfer to specific use cases with different cell types and limited training data.

## Method

### Data preprocessing

The prediction takes the target sequence as the input feature *X* and predicts the detailed editing outcomes *Y*, covering a diverse spectrum of events. The input sequence *X* was generally standardized, but the output events *Y* may have different annotation formats from method to method, depending on the coverage of the output events.

#### Transcode FORECasT data with the Lindel coding system

To convert FORECasT identifiers into higher-resolution Lindel classes, we built an editing outcome transcoding system based on FORECasT’s *indelmap* results, in which every unique indel was coupled with an identifier. We incorporated two indel labeling functions from separate studies and corroborated their labeling results to ensure the reliability of the conversion.

Our conversion pipeline relied on the CIGAR-string conversion function introduced by Liu et al. (Liu et al., 2022) and the IndelGen function from Chen et al. (Chen et al., 2019). The transcoding process took three inputs: an intact target sequence, a FORECasT identifier, and a paired editing outcome sequence. To map these inputs to the Lindel class, we followed these steps:

- We truncated the 79bp-long target sequence to 60bp leaving the cut site in the middle position (30bp).
- We used the IndelGen function to simulate all possible outcome-class label pairs for the truncated sequence.
- We matched the editing outcome sequence from the simulated outcomes and retrieved the corresponding Lindel class label.
- In parallel, we used Apindel’s CIGAR-string conversion function to map the identifier to the Lindel class.
- We outputted the Lindel class only when both Lindel classes were aligned.
- We aggregated the frequency vectors of all samples into a matrix with a dimension of (*n*_sample_, 557). The frequency matrix was used to construct AnnData. The name of each Lindel class, e.g.,” -1+1”, was stored in the var axis while the meta information about each guide, e.g., target sequence, parasite, and strand, was stored in the obs axis.

Overall, our transcoding system will turn a big data table in FORECasT format into a frequency matrix with a second dimension of 557.

We also considered an extension of Lindel’s 557 classes to 912 classes to improve the completeness of editing profiles. This coding system supported a maximum deletion length of 37bp. To convert FORECasT data to 912 classes, we skipped the target sequence truncation and ran *IndelGen* with the intact sequence and the cut site. Also, we broadened the criteria for valid deletion events in Apindel’s CIGAR-string conversion to include deletions from 1 to 37 bp. The insertion events of the 912 class coding system were the same as that of the 557 classes.

#### In-house embryonic data

For mouse embryonic editing data, we inputted the paired wild-type file, experimental file(s) of each gRNA to the Sanger analysis tools, Decodr (Bloh et al., 2021), to analyze gene editing efficiency with Sanger sequencing traces. We obtained each deletion sequence and its ratio directly from the analysis results. Given the software only provided the total 1 insertion ratio and the total 2 insertion ratio, we did further processing on insertion results. The 1 insertion event ratio was split by the estimated relative contribution of the inserted base. For 2 insertions, we just used the most frequent base pairs event with the ratio setting to be the sum of 2 insertion events. Each analyzed editing result was then mapped to the corresponding Identifier by our transcoding system based on the final editing sequence. A small portion of the files failed to be analyzed and were therefore discarded. We also did some manual filtering:

- Based on the output r-squared values, experimental files with poor fitting (r-squared < 0.75) were excluded.
- Experimental files with less than 10% effective Identifiers, indicating that the sum of events not encoded in our system was more than 90%, were filtered out.
- To mostly avoid misleading deconvolution results analyzed by Decodr, we also obtained analysis results from a second-party tool, Synthego, and verified the two output indel length event ratio vectors by setting the cosine similarity threshold to 0.8. Files with significant differences between the two editing tools were not included in our dataset.

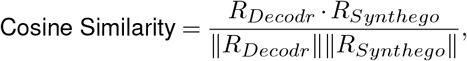

*Decodr*_*x*_ represents event vectors analyzed by Decodr with x indel length events, *Synthego*_*y*_ was y indel length events vector derived from Synthego. The length of the vector *R*_*Decodr*_, *R*_*Synthego*_ equal to the event number of the union of *Decodr*_*x*_ and *Synthego*_*y*_. The value of *R*_*Decodr*_ and *R*_*Synthego*_ vector element was the event ratio analyzed by the respective tool, with undetected events being assigned a value of 0.

Due to the limited size of sheep embryonic editing data, we simply aggregated the analysis results of experimental files by Decodr together for each gRNA, and using our Identifier to label the editing events, no further filtering were implemented.

Subsequently, the remaining results for the same gRNA were aggregated for single-clonal data, while for bulk data, the ratios were adjusted by multiplying them by the total number of embryos used in the experiment.

### Algorithmic design of *inDecay*

#### detection of microhomology events

To compile the data *x*_*i*_, we first needed to label whether an event was associated with microhomology-mediated end joining (MMEJ) or non-homologous end joining (NHEJ). In this study, we used a pairwise sequence identity matrix to identify all possible microhomologies within our samples. The process involved several steps, as follows.

We started with a target sequence and truncated it if it exceeded 79bp. At the designated cut site, we split the sequence into two fragments at the cut site. We then constructed a matrix where the left fragments occupied the rows (sequences before the cut site) and the right fragments filled the columns (sequences after the cut site). Each element in this matrix reflected the pairwise identity between nucleotides from the row and column fragments. We assigned a value of 1 for identical nucleotides and -1 for non-identical ones. Additionally, we can put 1 for the identical one and set 0 as the penalty for mismatched nucleotides. This setting will encourage the convolution filter to tolerate 1bp mismatch existing in the MH-track. In theory, more sophisticated scoring methods could be employed to differentiate between purines and pyrimidines.

To detect the diagonal signal patterns indicating microhomologies, we applied kernel convolution, using a 2×2 identity matrix as the convolution kernel. We selected diagonal patterns that exceeded a predefined threshold. In this study, we set the threshold to 3 as the default value in our package, meaning that only MH-tracks longer than 3bp were considered to induce MH deletion events. Each diagonal line in the matrix allowed us to identify its corresponding deletion start site and deletion length. For every filtered line, we denoted the coordinates of the head and tail ends as (*x*_1_, *y*_1_) and (*x*_2_, *y*_2_), respectively. This enabled us to assign the deletion start site as −1 × *y*_2_ and the deletion length as *y*_2_ + *x*_2_. The length of the microhomology was calculated using the formula 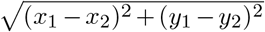. We mapped all filtered lines to their event labels and compiled them into a set *E*_*mh*_.

The detection process was encapsulated within a single-function model in our package, specifically in the function map.diagonal_map. For each target sequence paired with the guide *g*, this function allows us to obtain the specific set of microhomology events *E*_*g,mh*_ associated with *g*.

#### feature extraction

Our model, *inDecay*, was designed to be event-centered. Each event will be scored uniquely in the features except for features. Then we parameterized a regression function *f*_*p*_ to predict their frequency. We conceptualized deletion events as a competition between the decay of frequency and the recovery induced by microhomologies. Let *x*^(*i*)^ represent the features of a microhomology-mediated deletion event *i*, characterized by its deletion length *l*_*d*_, distance to the cut site *ss* and maximal microhomology length *l*_*mm*_. An MH deletion event can be firstly characterized using the following terms:

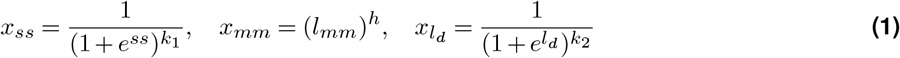

where *k*_1_ and *k*_2_ were the strength of decay with respect to deletion start site to (distance to the cuts ite) and deletion length respectively, *µ* and *σ* controlled the shape of decay concerning deletion length. *µ* was the lagging term only after which the decay happens. The feature of a Non-Homologous End Joining (NHEJ) deletion event can be obtained easily by setting the maximal microhomology length *l*_*mm*_ to 0. Alternatively, we can use the following transformation to extract features for the NHEJ deletion event if there is no lagging of decay in deletion length 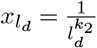 where *k*_2_ is the same in (1) which controls the strength of decay by deletion length.

When compiling the deletion features, we repeated each decay term with different hyperparameters to make the decay terms more stable. The full features extracted for deletion events include the raw start size *ss*, three *ss* decay terms with hyper-parameter *k*_1_ *0.5, *k*_1_, *k*_1_ *1.5, the raw deletion length *l*_*d*_, three *l*_*d*_ decay terms with hyper-parameter *k*_2_ * 0.5, *k*_2_, *k*2 * 1.5, the raw microhomology length and its power transformation with hyper-parameter *h*, the power transformation of *l*_*mm*_ allowing 1bp mismatch with hyper-parameter *h*, and whether the deletion was proximal or distal. In total, there were 13 event-specific features to describe deletion.

We modeled 1bp and 2bp insertions, a total of 20 insertion events, comprising four single nucleotide insertions, and 16 di-nucleotide insertions. For FORECasT’s coding system, several insertion events will be collapsed into one identifier. For our analysis, we only considered insertions that occurred at the cut site as valid. The nucleotide composition around the cut site largely influences the frequency of the insertion event. The full features we used to represent an insertion event were characterized by its insertion length, the existence of complementary nucleotides, the length of complements, the number of insertion indels collapsed in the identifier, and whether the complementary nucleotides were located left or right of the nucleotide. In total, there were 7 event specific features to describe insertion.

Additionally, we added several features that were shared among all events so that the model input could sense the global context of the target sequence. The shared features involved overall MH strength with/without mismatch, the prior probability of deletion and insertion ratio which were the output of ratio module *f*_*ratio*_, GC ratio of the guide RNA, which affected the resistance time of the CRISPR-gRNA complex Ma et al. (2016) and the one-hot encoding of sequence 9bp left to the PAM site (5 prime to 3 prime). The one-hot encoded features provide raw sequence information for the neural network to extract relevant hidden features. The two overall MH strengths were calculated from the filtered substitution matrixes *M*_*filter*_ obtained during the step of MH-detection. Distance matrix *D* denoted the distance of each point to the cut site. There were in total 41 shared features.

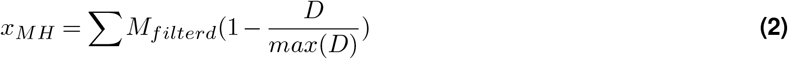

In summary, we ended up using 61 features (13 for del, 7 for ins, and 41 shared) as the input for the model.

#### Multistage model

*inDecay* model consisted of 2 modules, a ratio predictino module *f*_*ratio*_ and an event frequncy prediction module *f*_*p*_. The ratio predictor takes in the one-hot encoded sequences and predicts two scalers: the deletion ratio *r*_*del*_ and the insertion ratio *r*_*ins*_. A linear model was used, which was the same as Lindel (Chen et al., 2019). The backbone of *inDecay* can be flexible. One can use a linear regression model if preferring interpretability. In this study, we chose to use a multilayer perceptron (MLP) with only two hidden layers to parameterize the event predictor. The size of hidden variables was set to 128 and 64 to maintain a small model size. Additionally, we tested the backbone KAN proposed in (Liu et al., 2024), which stands for a new line of neural networks. KAN could reach a similar performance and enabled new functions like symbolic regression, but it ended up requiring a much larger parameter size and slowed the training.

A sample *g* with *n* possible events will have a feature matrix **X**_*g*_ ∈ ℝ^*n*\*61^. The rate of a event *z*_*i*_ was predicted by the module *f*_*p*_ from the features array *x*_*i*_, depending on whether *i* was an insertion or deletion event:

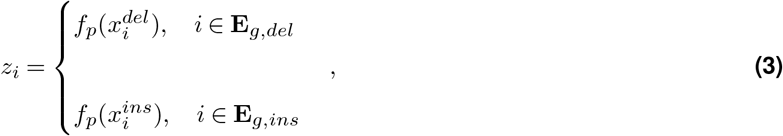

Finally, the full event probability matrix **P**_*g*_, **P**_*g*_ ∈ ℝ^*n*^ was obtained by transforming the rate into probability:

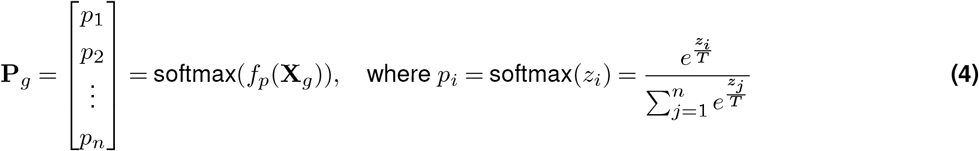

### Train test split

For somatic cell lines training, we split the entire dataset into train-val and test sets. The test set was first held out to benchmark different models, and the remaining samples acted as a train-val set. A train-val split was called with different random seeds for different training repeats to sample a unique training data set. Specifically for the retraining of FORECasT, we only included guides with over 1000 total count and belonging to the forward strand in the training process, which was required by (Allen et al., 2019). To find a set of samples whose editing profiles were well-represented across all cell lines, we selected test set samples with the following criteria: 1) the total count of the sample was greater than 1000 in all five cell lines; 2) Both the insertion ratio and deletion ratio were not 0, which was to remove insertion-free guides and deletion-free guides. Finally, 1133 guides passed the above criteria, and they were withheld during the training process.

For cross-cell line transfer learning, in each sample size setting, we randomly selected a certain number (*N*_*sample*_) of samples 10 times using different seeds in the mESC and iPSC datasets, respectively. And 20% of the samples (0.2*N*_*sample*_) were used for validation. The test set for each cell line was the same as the dataset used in somatic cell lines.

For few-shot learning in embryonic data, we used a leave-one-out strategy, with 1 gRNA for testing and the remaining data for training, 20% of which were used for validation.

### Model training

All the models were developed with PyTorch version 1.12.1. Our study employed two distinct loss functions in somatic cell line training. The first was a cross-entropy loss, which was used to optimize each event predictor model based on the observed event frequency and the predicted probability *P*. Both Lindel and FORECasT followed the same training schema. In addition, we introduced a multi-nomial loss function that computed the negative log-likelihood of observing the true event counts **n** with a multinomial distribution parameterized by the predicted probability *P*.

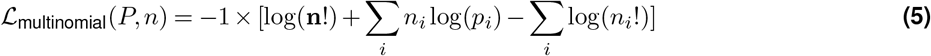

For few-shot learning in mouse embryonic editing data, we used a weighted cross-entropy loss penalizes predictions further from the true label *y*_*i*_ for gRNAs with larger embryo sample sizes *n*_*i*_, also with an additional term to regularize the trained weights *t* by minimizing its *l*_2_ − *norm* with initial pre-trained weights *w*. The default *λ* was set to be 0.3.

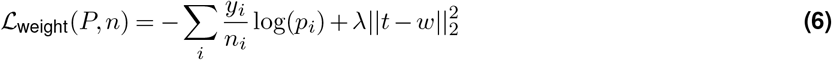

Also, in the last layer, we used the Temperature in the Softmax strategy to control the randomness of predictions by scaling the logits *p*_*i*_. The default temperature *T* was set to be 1 for somatic cell training and 1.4 for embryonic editing data fine-tuning.

*Adam* optimizer was employed with a learning rate of 0.0003 for both training and few-shot learning. The weight decay parameter of Adam was set to be 0.0001.

### Model evaluation

Multiple metrics were included in our study to evaluate the prediction performance. The KL divergence measures how one probability distribution differs from a second probability distribution,

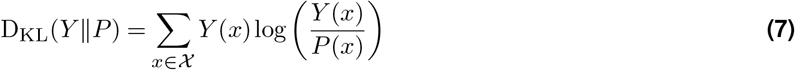

where *Y* (*x*) denoted the observed events distribution while *P* (*x*) was the predicted events distribution,

In addition, we introduced top-k precision to measure the accuracy of the most frequent events. These two metrics took the concept of precision@k and recall@k from the field of recommender systems, we referred to the metric as Top-k events recall and we normally chose k equals to 1, 5, and 10 to check the prediction precision in different levels in the paper.

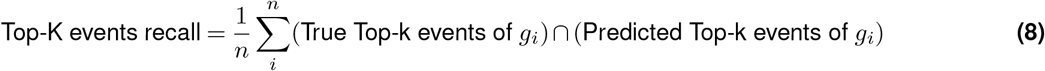

Besides, Kendall’s Tau is a correlation coefficient that measures the association between two ranked variables. It is a non-parametric statistic that evaluates the similarity of the orderings of the data.

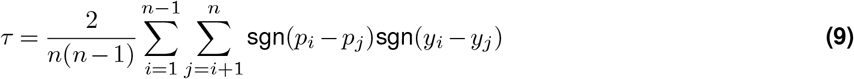

Finally, we employed r-square to measure the concordance of a wide range of ratios, including frameshift proportion and deletion-insertion ratio

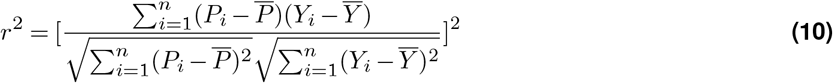

where *P*_*i*_ was the predicted frameshift or deletion ratio vectors for *g*_*i*_, while *Y*_*i*_ was the observed frameshift or deletion ratio vectors for *g*_*i*_.

All metrics above, except those associated with r-square, were calculated with two probability distributions. It’s, therefore, possible to reuse these metrics after transforming the event probability distribution into an indel length distribution. Despite a lower resolution than the event-based distribution, transforming into an indel length distribution enables the comparison of the results of different methods regardless of the coding system of use.

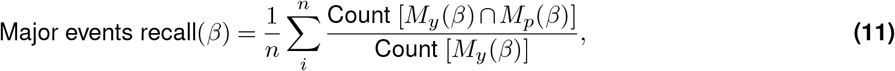

where *M*_*y*_(*β*) were the events that have a true ratio larger than the threshold *β, M*_*p*_(*β*) were the events that predicted ratio larger than the threshold *β*, and *n* was the number of available gRNAs. This metric will be omitted for the gRNA if Count (*M*_*y*_(*β*)) = 0.

## Data Availability

Self-targeting data is available for download at https://figshare.com/articles/processed_mutational_profiles/7312067. The transformed self-targerting data and sanger sequencing data that are ready to reproduce the *inDecay* can be accessed via the data download script located at https://github.com/StatBiomed/inDecay. Decoded data tables are included in the supplementary materials.

## Code Availability

inDecay is implemented as a standard Python package and the source codes are freely available at https://github.com/StatBiomed/inDecay. All analysis notebooks are also included in the repository for reproducibility.

## Author Contribution

Y.H., G.L., P.L., C.C. conceived the project. W.Z. designed the inDecay model and performed the benchmarking part with help of L.Y. and Y.H. L.Y. pre-processed the in-house embryonic editing data and compiled the dataset with support from W.Z., G.L., Y.H. and C.C. L.Y. and W.Z. performed all the transfer learning parts with assistance of Y.H. G.L. designed and organized all the embryonic editing experiments. G.W. and K.H. provided mouse embryonic editing data; S.C. generated the bovine, porcine, goat, rabbit embryonic editing data; J.S. and Z.L. performed the rabbit embryonic editing experiments; X.L. and M.W. performed the porcine embryonic editing experiments. H.W. provided resources for rabbit, porcine and goat embryonic editing. G.L. controlled the quality of the original editing data. P.L. provided resource for the project. W.Z., L.Y. and Y.H. wrote the manuscript from the input of all authors.

## Supplementary Information

**Figure S1.**
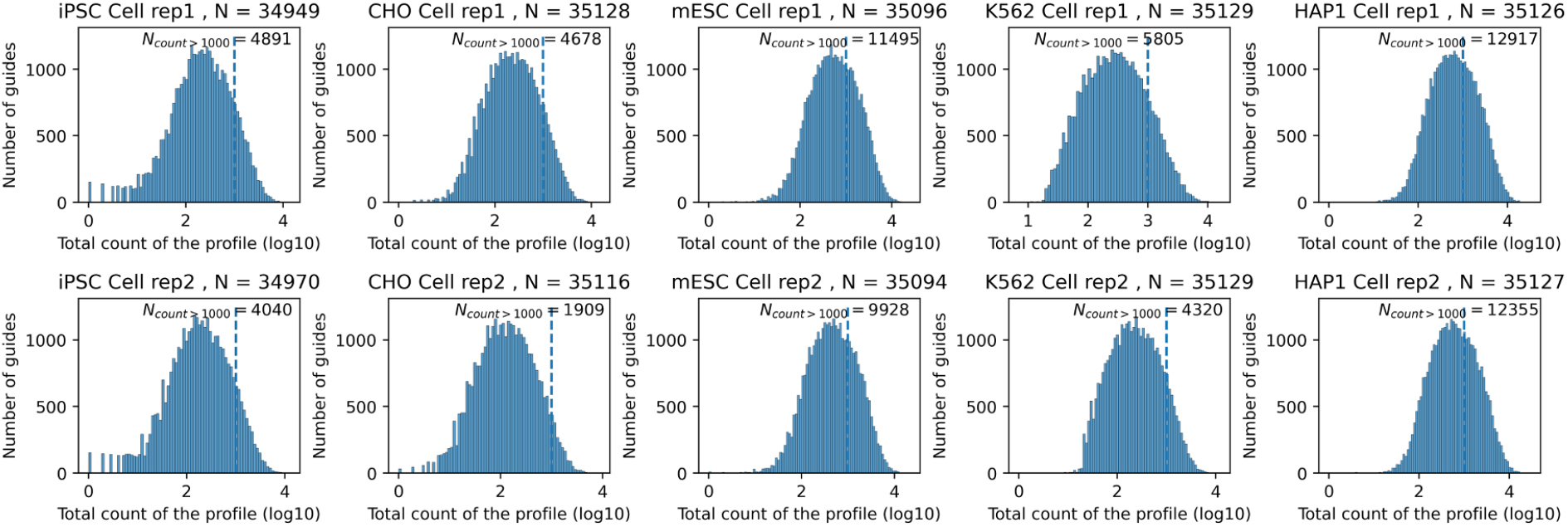
Total read count distribution of FORECasT’s multi-cell-line dataset. This figure shows the distribution of total read counts in FORECasT’s multi-cell-line dataset. Each panel displays the density of target sequences based on their total number of edited reads. The x-axis uses a log10 scale, with a dashed line marking the 1,000 count position. Each cell line includes two repeats, with the total sample size mentioned in the subtitle. The number of samples exceeding 1,000 reads is also noted for each cell line and experimental repeat.

**Figure S2.**
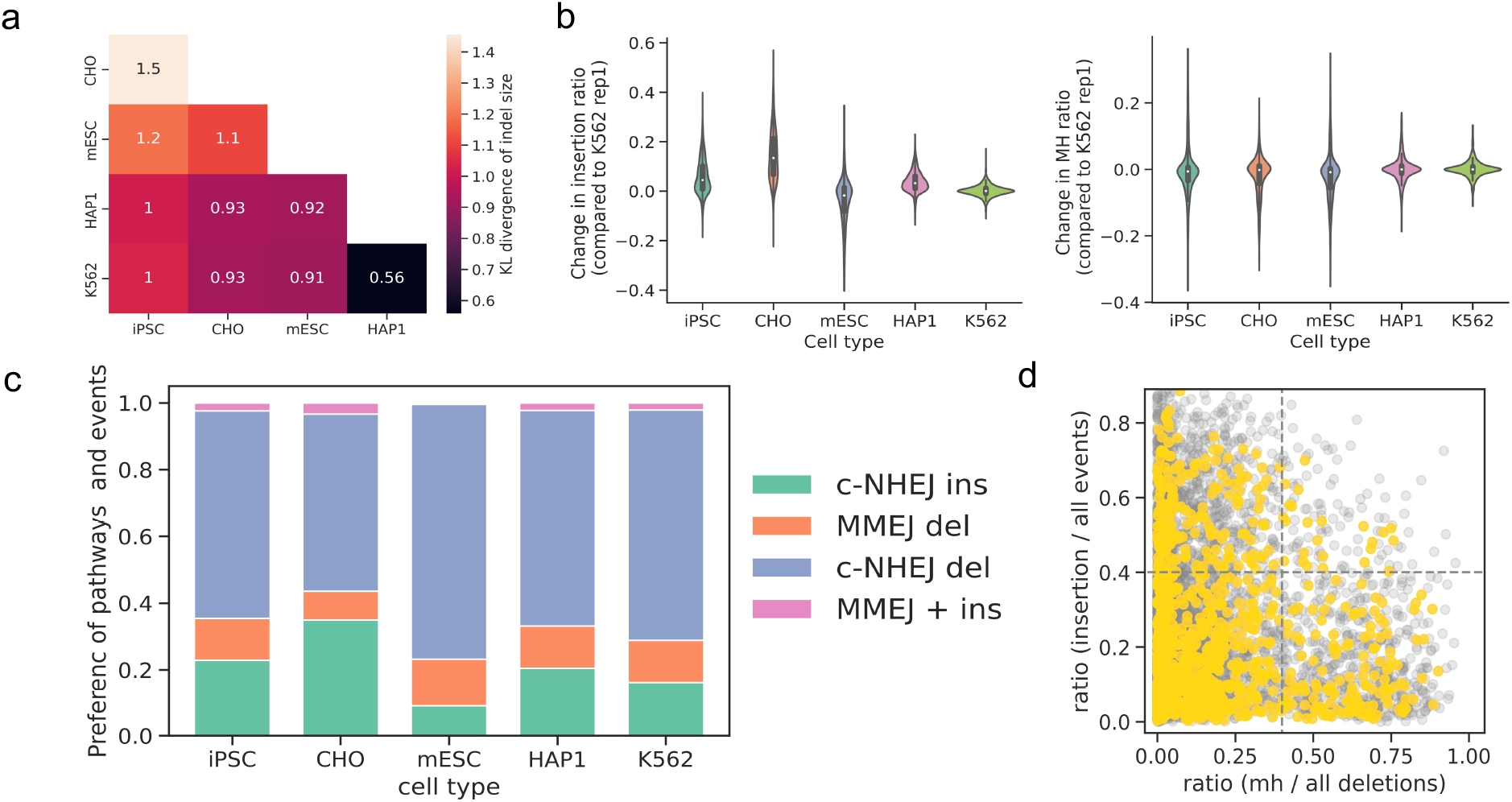
Cell type variability of editing profiles. (a) The pairwise KL divergence of indel-size distribution. (b) Violin plot showing the changes in insertion ratio (left) and MH strength (right). The ratio changes were calculated relative to replicate 1 of K562 cells between matched guides (replicate 2 to replicate 1 for K562). (c) The proportion of samples dominated by each repair pathway. In the stacked bar plot, color showed different repair pathways. Guides with both MH ratio and insertion ratio higher than 0.4 are noted with “MMEJ + ins”. (d) Scatter plot showing MH ratio (x-axis) and the ratio ratio (y-axis) of each guide from K562 cells. The gray dots are background generated by pooling samples from all cell lines.

**Figure S3.**
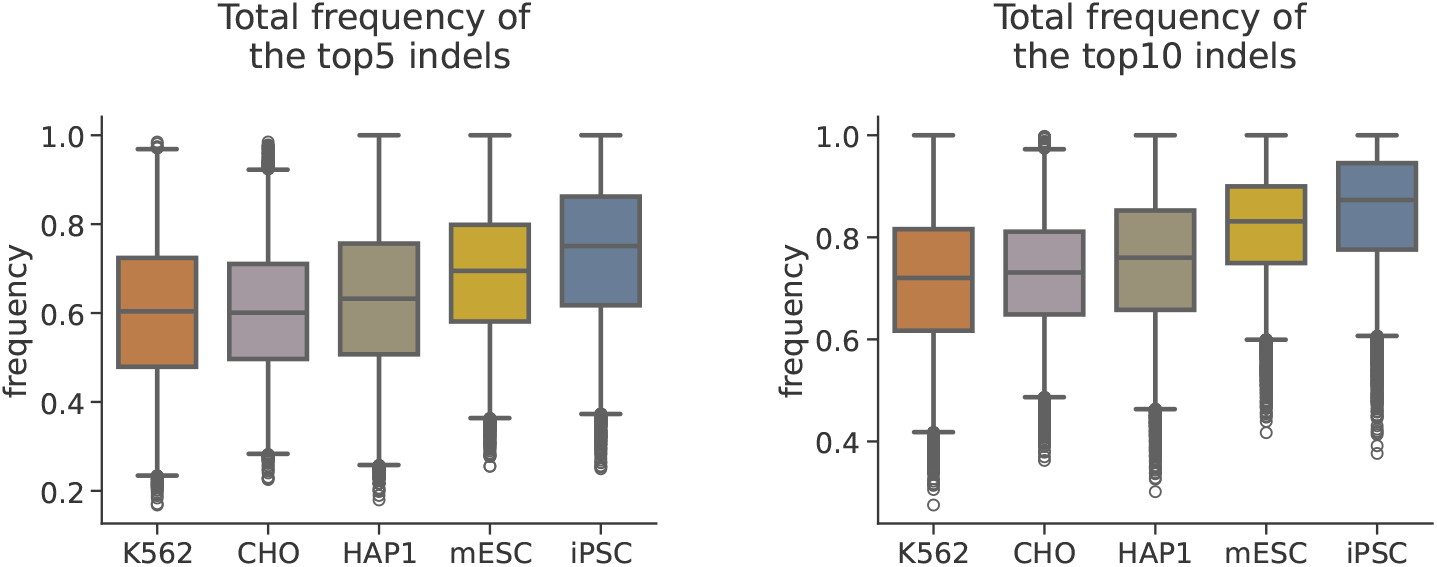
The frequency of the most frequent indels. Box plots showing the total frequency of the top 5 and top 10 indels across different cell lines (K562 n=10174, CHO n=8575, HAP1 n=17225, mESC n=15532, iPSC n=7831). The left plot represents the top 5 indels, and the right plot represents the top 10 indels. Each box plot uses a whisker length of 1.

**Figure S4.**
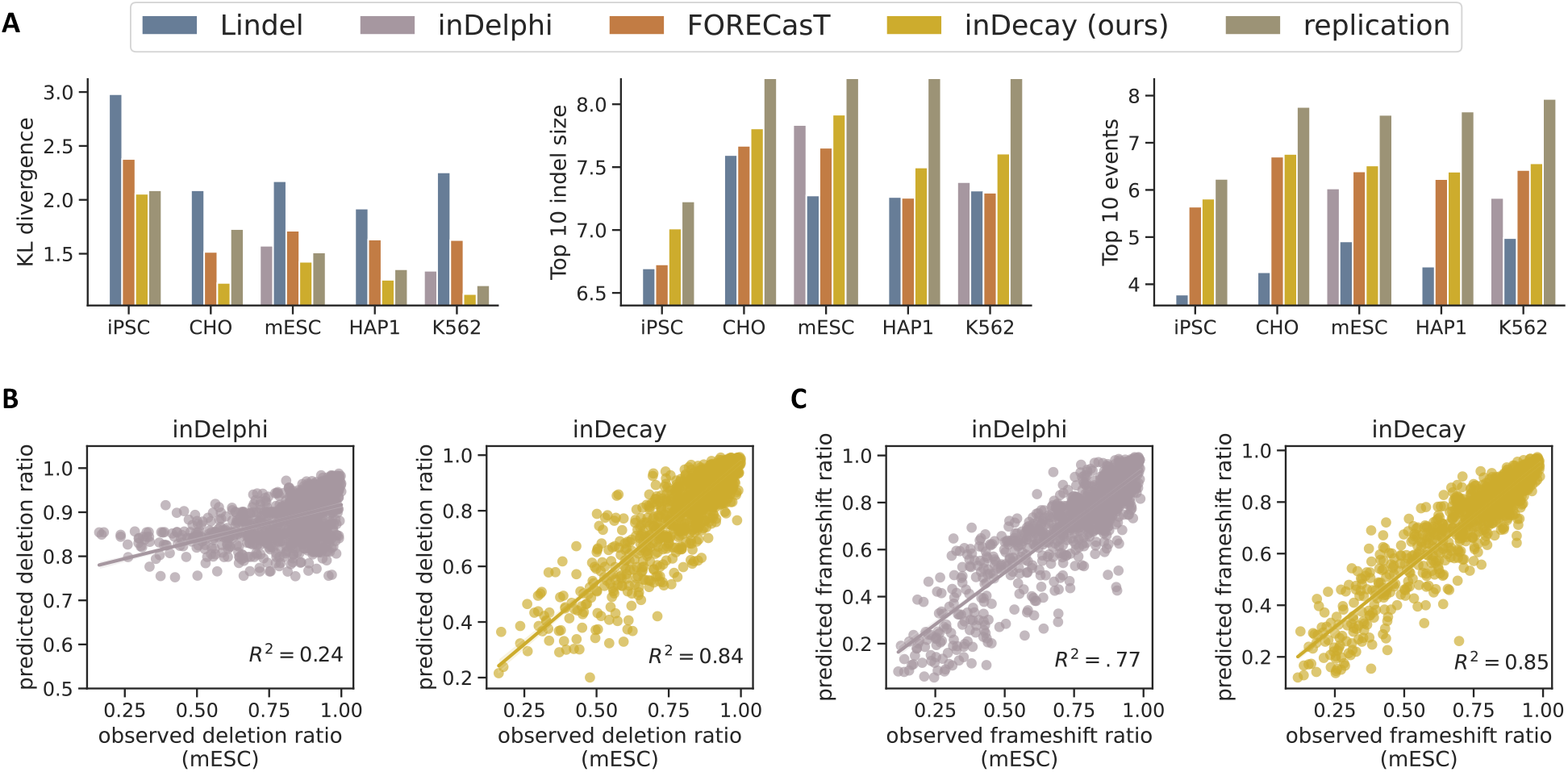
Comparison of inDecay with other method and ratios. **(A)** Three additional evaluation metrics for benchmarking different repair outcome prediction methods. The left panel shows the KL divergence between the observed and predicted outcome probabilities. The middle and right panels show the recall of the top 10 indel sizes and top 10 events. These methods are trained, evaluated, and visualized in the same way as in Figure 2. (**B**) The deletion ratio for each test set sample is shown, compared with predicted ratios from inDelphi (gray) and our inDecay (yellow). The *R*^2^ score is calculated and also presented as a bar plot in Figure 2. (**C**) Scatter plots illustrate the frameshift ratio of each sample, using the same color scheme as in (B).

**Figure S5.**
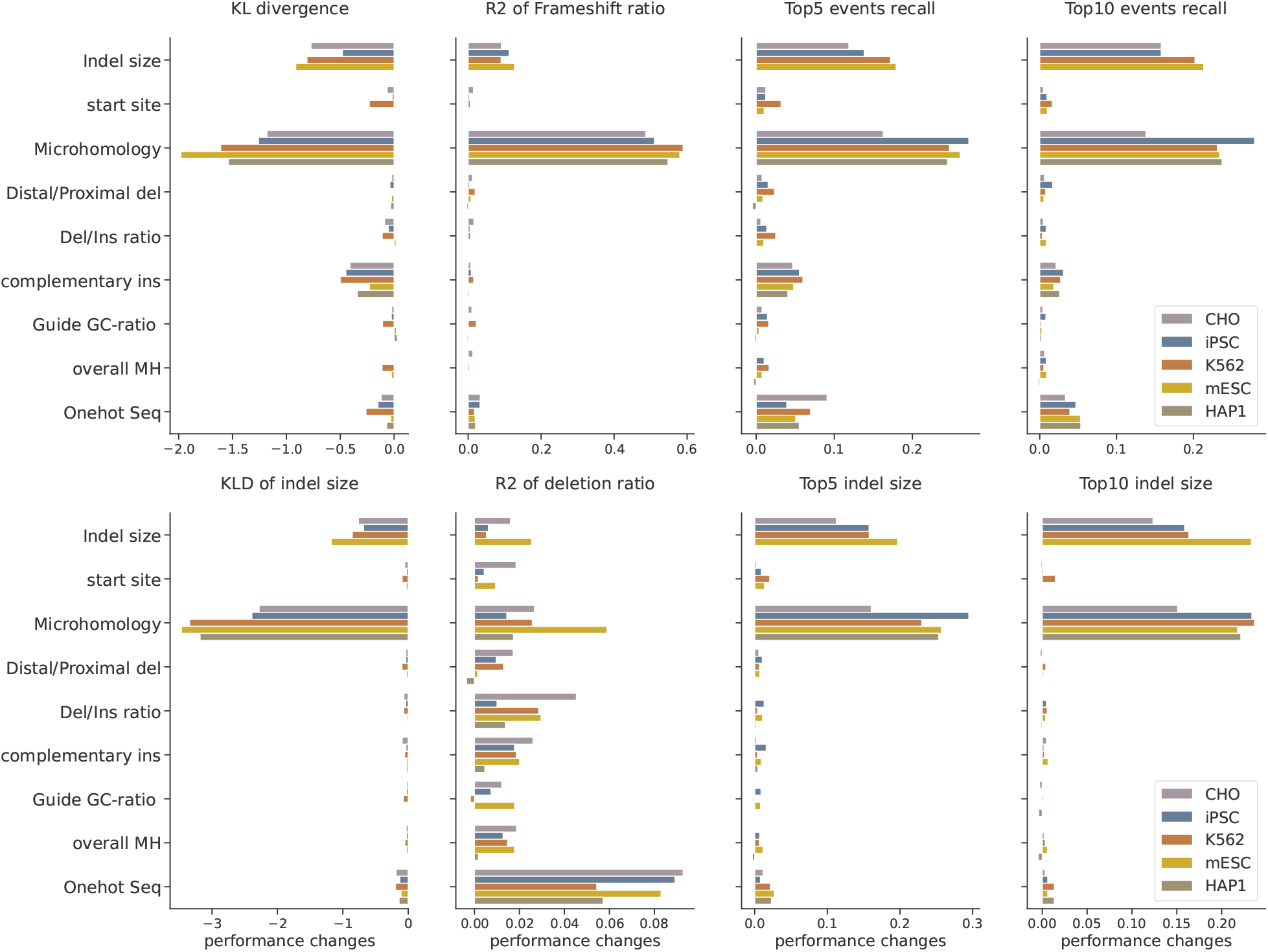
Feature importance of the ablation study. In our ablation study, we retrained the inDecay model, selectively masking certain input dimensions from the same feature set. Each panel illustrates the evaluation metrics used to calculate feature importance. The Y-axis indicates the feature set being masked, while the X-axis shows the performance difference, which was obtained by the ratio of difference (the original model minus the masked model) over the original value. Except for the two KL divergence, higher value indicates. The color of the bar denotes the cell-line of the training data.

**Figure S6.**
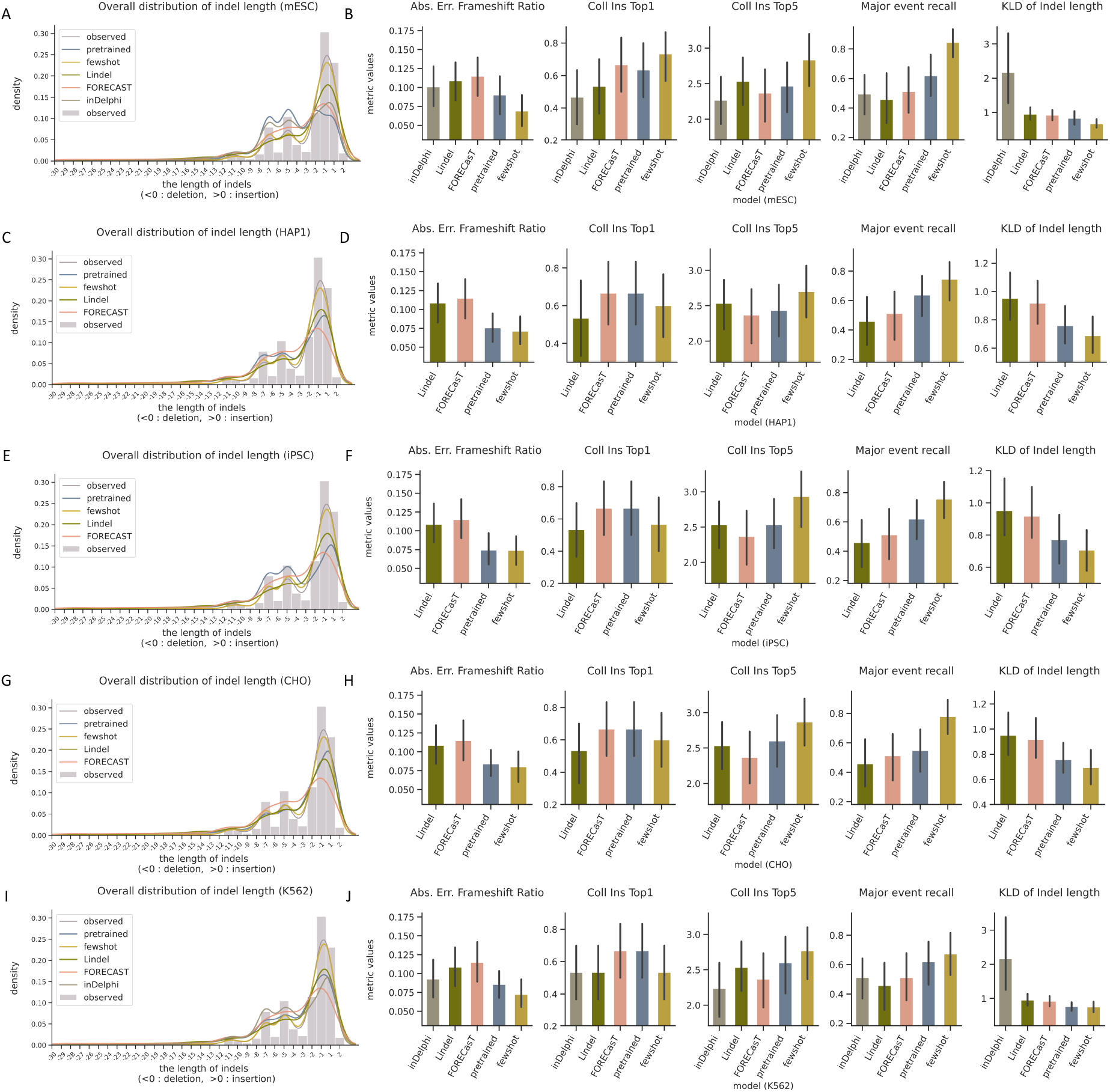
Transferring and evaluating inDecay and other methods in our collected embryonic editing dataset based on 5 somatic cell lines. (**A, C, E, G, I**) The overall distribution of indel length events, inDecay pretrained and few-shot learning model pre-trained on mESC, HAP1, iPSC, CHO and K562 cells respectively. (**B, D, F, H, J**) Evaluate the performance of each tool/ setting by absolute error of frameshift ratio, Top1 events recall (merging same insertion length events), Top5 events recall (merging same insertion length events), Major event recall (*β* = 0.2) and KL Divergence of indel length events, with inDecay base model listed on the label of x-axis.

