## Supplementary Information for "Accurate prediction of CRISPR editing outcomes in somatic cell lines and zygote with few-shot learning"

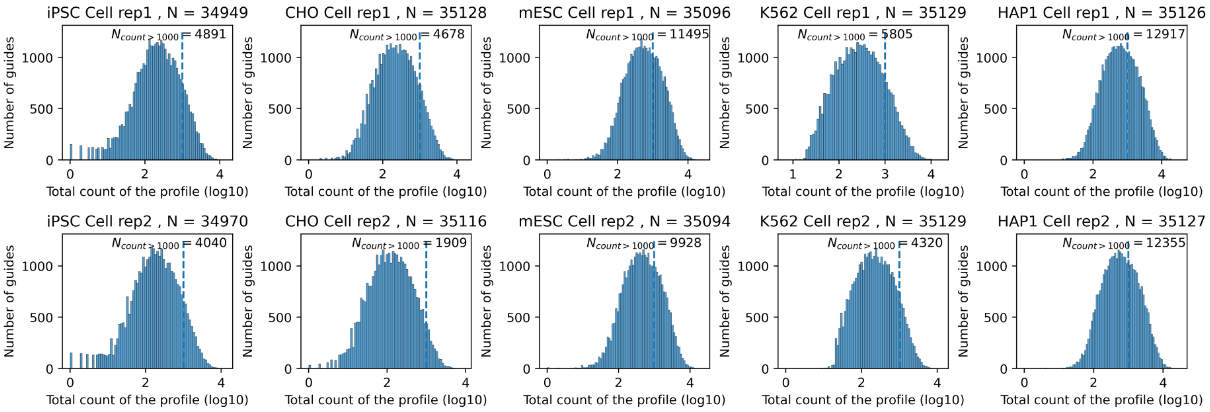


**Figure S1. Total read count distribution of FORECasT's multi-cell-line dataset.**
This figure shows the distribution of total read counts in FORECasT's multi-cell-line dataset. Each panel displays the density of target sequences based on their total number of edited reads. The x-axis uses a log10 scale, with a dashed line marking the 1,000 count position. Each cell line includes two repeats, with the total sample size mentioned in the subtitle. The number of samples exceeding 1,000 reads is also noted for each cell line and experimental repeat.

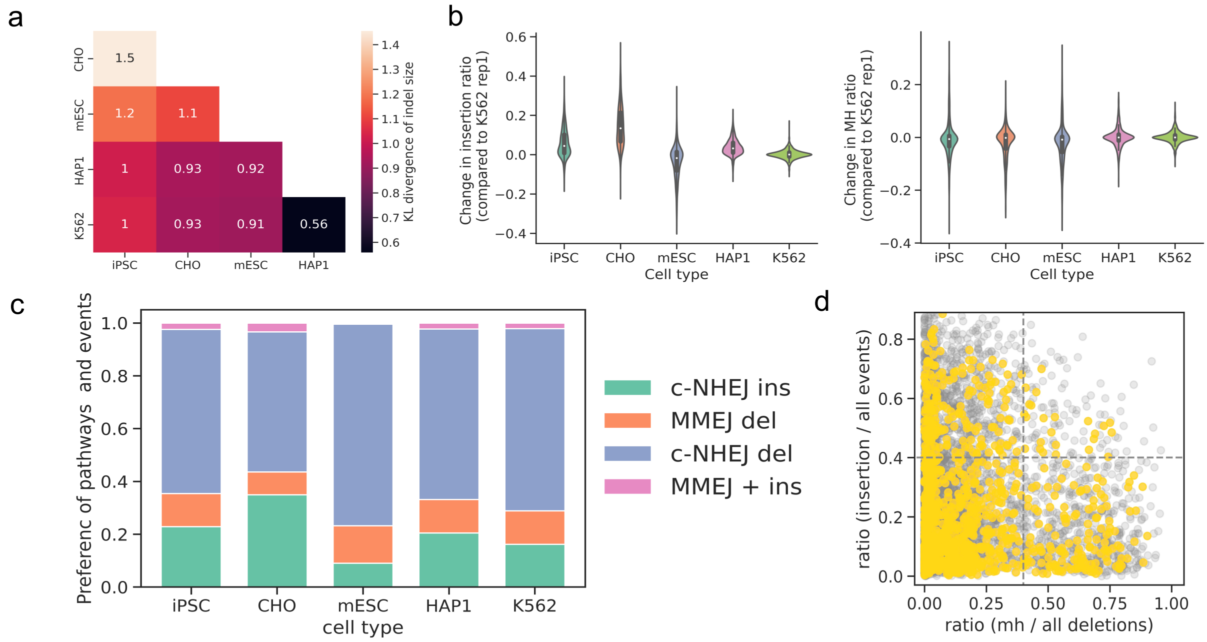


**Figure S2. Cell type variability of editing profiles.**
(a) The pairwise KL divergence of indel-size distribution. (b) Violin plot showing the changes in insertion ratio (left) and MH strength (right). The ratio changes were calculated relative to replicate 1 of K562 cells between matched guides (replicate 2 to replicate 1 for K562). (c) The proportion of samples dominated by each repair pathway. In the stacked bar plot, color showed different repair pathways. Guides with both MH ratio and insertion ratio higher than 0.4 are noted with "MMEJ + ins". (d) Scatter plot showing MH ratio (x-axis) and the ratio ratio (y-axis) of each guide from K562 cells. The gray dots are background generated by pooling samples from all cell lines.

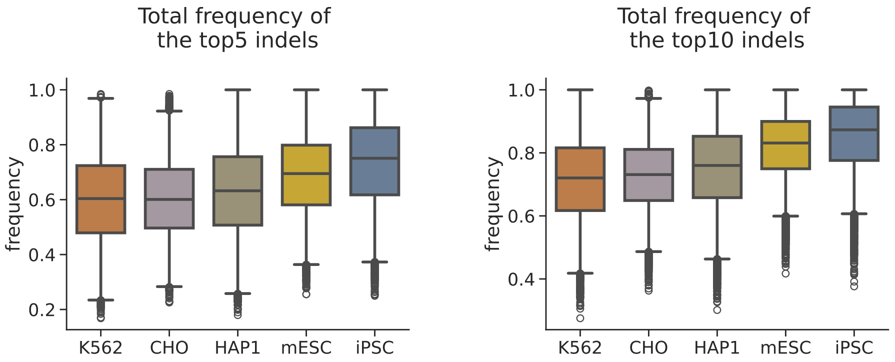


**Figure S3. The frequency of the most frequent indels**
Box plots showing the total frequency of the top 5 and top 10 indels across different cell lines (K562 n=10174, CHO n=8575, HAP1 n=17225, mESC n=15532, iPSC n=7831). The left plot represents the top 5 indels, and the right plot represents the top 10 indels. Each box plot uses a whisker length of 1.

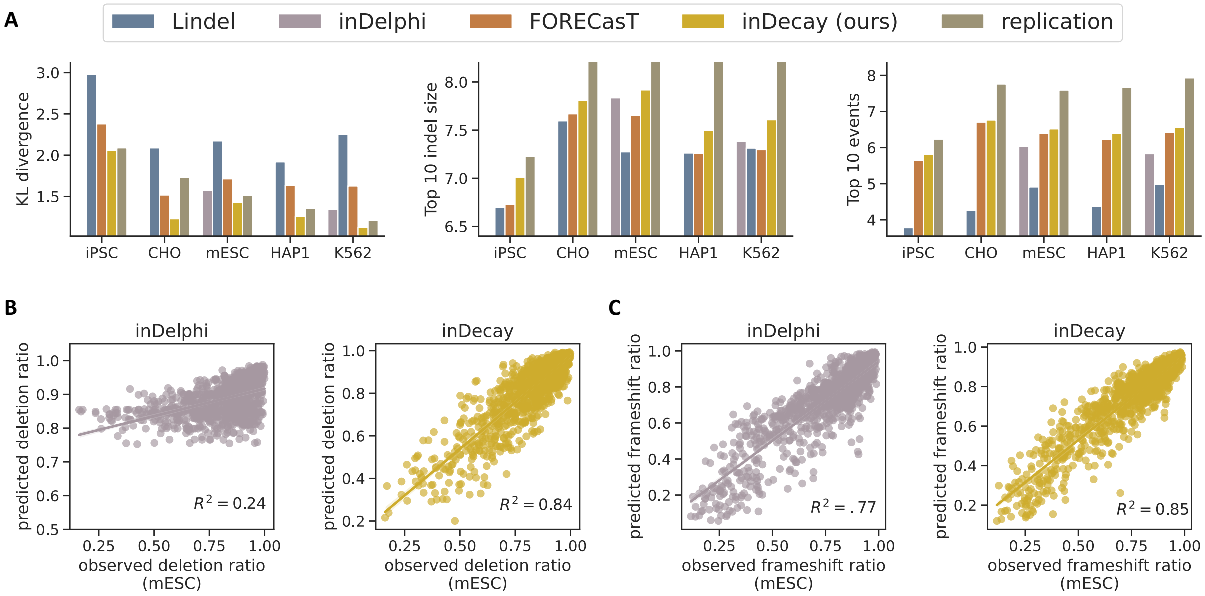


**Figure S4. Comparison of inDecay with other method and ratios**
(**A**) Three additional evaluation metrics for benchmarking different repair outcome prediction methods. The left panel shows the KL divergence between the observed and predicted outcome probabilities. The middle and right panels show the recall of the top 10 indel sizes and top 10 events. These methods are trained, evaluated, and visualized in the same way as in Figure 2. (**B**) The deletion ratio for each test set sample is shown, compared with predicted ratios from inDelphi (gray) and our inDecay (yellow). The $R^2$ score is calculated and also presented as a bar plot in Figure 2. (**C**) Scatter plots illustrate the frameshift ratio of each sample, using the same color scheme as in (B).

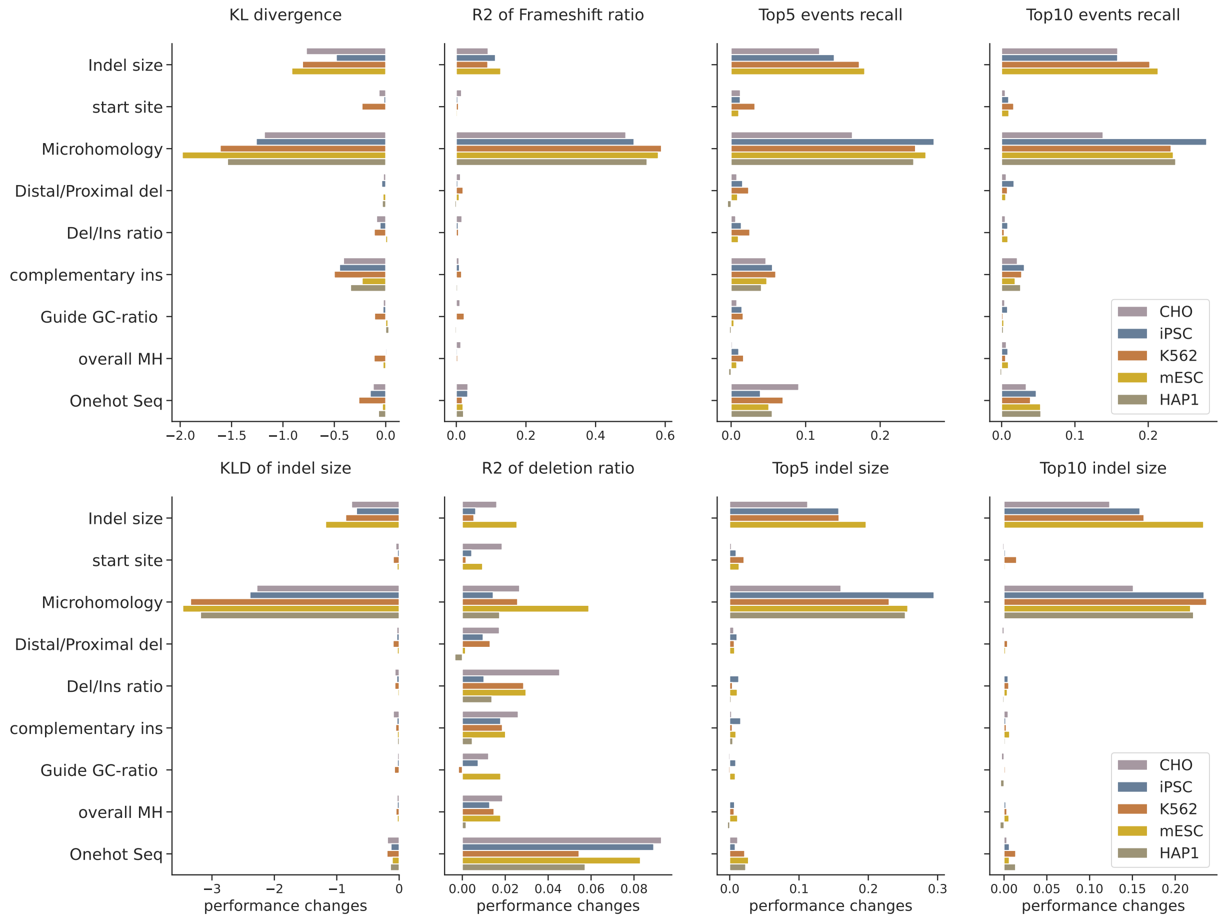


**Figure S5. Feature importance of the ablation study**

In our ablation study, we retrained the inDecay model, selectively masking certain input dimensions from the same feature set. Each panel illustrates the evaluation metrics used to calculate feature importance. The Y-axis indicates the feature set being masked, while the X-axis shows the performance difference, which was obtained by the ratio of difference (the original model minus the masked model) over the original value. Except for the two KL divergence, higher value indicates . The color of the bar denotes the cell-line of the training data.

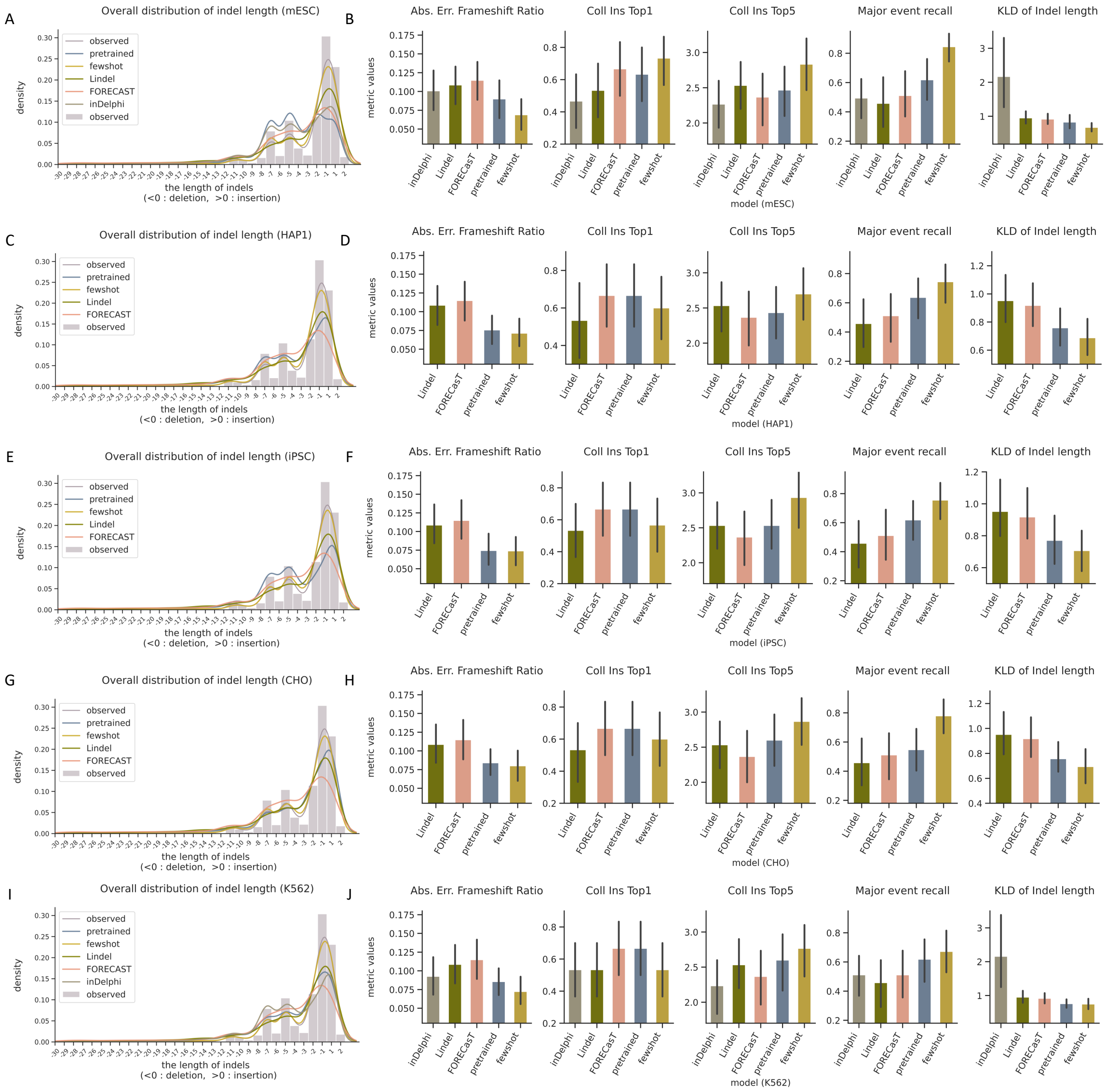


**Figure S6. Transferring and evaluating inDecay and other methods in our collected embryonic editing dataset based on 5 somatic cell lines**

(**A, C, E, G, I**) The overall distribution of indel length events, inDecay pretrained and few-shot learning model pre-trained on mESC, HAP1, iPSC, CHO and K562 cells respectively. (**B, D, F, H, J**) Evaluate the performance of each tool/ setting by absolute error of frameshift ratio, Top1 events recall (merging same insertion length events), Top5 events recall (merging same insertion length events), Major event recall ($\beta$ = 0.2) and KL Divergence of indel length events, with inDecay base model listed on the label of x-axis.

**Table 1. The guide sequence of designed gRNA and total embryo samples used**

| Item | Species | Guide | Total Embryos |
|---|---|---|---|
| Adam6 | Mouse | ACTTGGCGTTACATCTCAT | 27 |
| Adgre1 | Mouse | AATATATTTAAAAGCAAGAA | 42 |
| Ctnnb1 | Mouse | TTTAGCAGTTTTGTCAGCTC | 52 |
| Dcbld2 | Mouse | ACAAGTACATACCTGAACTC | 30 |
| DIRK | Mouse | GACAGATGGGGGTGTCGTTT | 30 |
| ELF5 | Mouse | TCTGATGGACACCGGACACC | 21 |
| EPHA2 | Mouse | TGAAATAGCCTTCTTCACAC | 4 |
| H2AB1-sg1 | Mouse | CGAGTACTGGAACAGCCAGC | 36 |
| H2AB1-sg2 | Mouse | AGGCAAAGGGCAGAGGGCAG | 11 |
| H2D1-sg1 | Mouse | GGGGCTCCTCGAGGCCGGGC | 7 |
| H2D1-sg2 | Mouse | GAACCACTGCTCTTGGCCCT | 28 |
| IL2rg | Mouse | GAGCAGCTGAAGGACTAAGA | 7 |
| IL5 | Mouse | TTCTGACTCTCAGCTGTGTC | 38 |
| IL6 | Mouse | TCCTCTCTGCAAGTAAGTGA | 16 |
| IL7 | Mouse | CTCCCGCAGACCATGTTCCA | 38 |
| Kat2a | Mouse | CTGCCTTAACTACTGGAAGC | 57 |
| Mettl14-sg1 | Mouse | ACATCCCTGATGAAATTCTG | 71 |
| Mettl14-sg2 | Mouse | AAATAGCAAAGATGAACAGA | 28 |
| mHV5-sg7 | Mouse | TGTGGGGTTCAGGTGGCCTA | 34 |
| Mrgpra | Mouse | CCAATGGACGAAACCCTCCC | 9 |
| MTAP | Mouse | GCGTTTGAAGCTACAGCTCA | 44 |
| PDL1 | Mouse | GACGTCAAGCTGCAGGACGC | 25 |
| Rag1 | Mouse | TGTGTGGGGGTGCCACTCCA | 9 |
| Rag1-sg1-4 | Mouse | TCATGCAAGGCAGGGGCTCC | 9 |
| sry | Mouse | TTTGCATGCTGGGATTCTGC | 66 |
| stk4-sg2 | Mouse | GTTTTGGAGATGATTGGAGA | 33 |
| stk4-sg3 | Mouse | CAGTTTGGGGATGAACTTCA | 7 |
| GM-sg1 | Mouse | ACTTCACCAAATAAAGAGAA | 35 |
| GM-sg2 | Mouse | ATGACTTCTTAATAGGTTCC | 42 |
| Sorl1 | Mouse | TCTCTAGCAGTGCTGGAGCC | 97 |
| SOCS2-sg83-F | Sheep | GTCTTAACAGATATTGTTAGT | 14 |
| SOCS2-sg96-F | Sheep | GTATTCGATGCGAAGATTAGT | 14 |
